# Relevance of the synergy of surveillance and populational networks in understanding the Usutu virus outbreak within common blackbirds (*Turdus merula*) in Metropolitan France, 2018

**DOI:** 10.1101/2024.07.22.604715

**Authors:** Malika Bouchez-Zacria, Clément Calenge, Alexandre Villers, Sylvie Lecollinet, Gaelle Gonzalez, Benoit Quintard, Antoine Leclerc, Florence Baurier, Marie-Claire Paty, Éva Faure, Cyril Eraud, Anouk Decors

## Abstract

Usutu virus (USUV) was first isolated in Africa in 1959 and has since spread to and through Europe with a typical enzootic mosquito-bird cycle. In France, it was first detected in birds in 2015, but during summer 2018 the spread of USUV was particularly significant throughout the country, killing mainly common blackbirds (*Turdus merula*) and to a lesser extent great grey owls (*Strix nebulosa*), among other captive and non-captive wild bird species. Previous studies of USUV in France have focused on reconstructing pathways of introduction, but not on structural aspects of virus spread within the country. Data (RT-PCR of geolocated dead birds) on this 2018 outbreak were collected through both an event-based wildlife network named SAGIR and the health surveillance of the French-speaking Association of Zoo Veterinarians (AFVPZ). In addition, common blackbird populations could be monitored through another network (REZOP). Statistical analysis (spatial, temporal, spatiotemporal and environmental determinants) of the SAGIR and AFVPZ network data helped to highlight the early appearance of separate large clusters of USUV cases in mid-July 2018, the subsequent diffusion into smaller and secondary clusters at the end of August 2018, and a meanwhile enlargement of the first clusters with an increase in the number of cases. High human density (top 10.5% densest areas in France) and wetland concentration (top 19.3% most likely wetland areas) were significant factors in USUV case locations. Using generalised additive mixed models on REZOP data, we also highlighted the decline in common blackbird population trends in areas with medium and even more with high USUV pressure (areas defined based on SAGIR-AFVPZ data) following the 2018 outbreak (respectively −7.4% [−11.4; −3.9]_95%_ and −15.7% [−16.2; −9.1]_95%_). A large area (radius ∼150 km) in the centre and centre-west of France, and smaller areas in the south-east, north and north-east of France (each with a radius ∼ 50 km) were particularly affected. We conclude on the importance to work with synergistic networks to assess infection spread in wild bird species, as well as the negative impact of an emerging arbovirus. The responsiveness of such a network system could be improved by automating alerts.

## Introduction

The Usutu virus (USUV), so called because of its isolation in *Culex neavei* mosquitoes in 1959 (McIntosh 1985) near the Usutu river in Swaziland (Eswatini since 2018) is an arbovirus of the *Flaviviridae* family, *Orthoflavivirus* genus (Clé et al. 2019), belonging to the Japanese encephalitis virus serocomplex, and phylogenetically close to Japanese Encephalitis Virus (JEV) and West Nile Virus (WNV) (Calisher and Gould 2003). After South Africa, the virus was detected in other African countries: Central African Republic, Senegal, Ivory Coast, Nigeria (Nikolay et al. 2011), Uganda (Nikolay et al. 2011; Mossel et al. 2017), Burkina Faso (Nikolay et al. 2011), Mali, Madagascar (Chevalier et al. 2020), Kenya (Ochieng et al. 2013), Tunisia (Ben Hassine et al. 2014; M’ghirbi et al. 2023), Morocco (Durand et al. 2016), and Israel (Mannasse et al. 2017). The first known occurrence in Europe dates back to 1996 in Italy (a retrospective finding in dead birds, mainly blackbirds) (Weissenböck et al. 2013). Since then, first detections of USUV followed in other European countries: Austria (Weissenböck et al. 2002) in 2001, Czech Republic (Hubálek et al. 2008a)) in 2004, Hungary (Bakonyi et al. 2007) in 2005, Poland (Hubálek et al. 2008b), Spain (Busquets et al. 2008) and Switzerland (Steinmetz et al. 2011) in 2006, Serbia in 2009 (Lupulovic et al. 2011), Germany (Jöst et al. 2011), Greece (Chaintoutis et al. 2014) and Slovakia (Csank et al. 2018) in 2010, Croatia (Barbic et al. 2013) in 2011, Belgium (Garigliany et al. 2014) in 2012, France (Lecollinet et al. 2016) in 2015, Netherlands (Rijks et al. 2016) in 2016 and United Kingdom (Folly et al. 2020) in 2020 (Vilibic-Cavlek et al. 2020).

The analysis of 92 complete USUV genomes, including 77 genomes from mosquito, bird and bat species of Germany (2010-2014) highlighted that i) USUV can be classified in different lineages (Engel et al. 2016), (eight in number – three African and five European (Cadar et al. 2017; Clé et al. 2019)), ii) the most common ancestor emerged in Africa at the beginning of the 16^th^ century, iii) USUV was regularly introduced from Africa during the last 75 years (a first introduction from 1950 through 1960’s in Western Europe (Spain), a second one between 1970 and the 1980’s in Central Europe (Austria) and a third one around 1996 in Western Europe (Spain)) and iv) *in situ* evolution could explain the genetic diversity of European lineages, while extensive gene flow drove African ones. In addition, flyway networks of migratory birds were strongly consistent with spatial movements observed through genetic data and suggested the possibility of USUV exportation from Africa i) to Spain via the east Atlantic and/or Black Sea/Mediterranean flyways and ii) to Central Europe through the Black Sea/Mediterranean flyway (Engel et al. 2016).

The natural life cycle of USUV involves passeriform and strigiform birds as amplifying hosts and ornithophilic species of mosquitoes as vectors (Clé et al. 2020). The virus has been detected in several Culicidae (*Culex*, *Aedes, Culiseta*, *Mansonia*) (Clé et al. 2019) but *Cx. pipiens* (experimentally shown to be USUV competent (Martinet et al. 2023)) is considered as the main vector in Europe (Becker et al. 2012; Martinet et al. 2023). To date, several incident or dead-end hosts (which certainly develop a low viremia, insufficient to re-infect mosquitoes (Martinet et al. 2023)) have been identified (Vilibic-Cavlek et al. 2020): horses (Durand et al. 2016; Bażanów et al. 2018; Csank et al. 2018), dogs (Durand et al. 2016), rodents (Diagne et al. 2019), squirrels (Romeo et al. 2018), wild boar (Escribano-Romero et al. 2015; Bournez et al. 2019) and roe deer (Bournez et al. 2019). The virus has also been detected in bats, whose epidemiological role is as yet undetermined (Cadar et al. 2014). The virus has a zoonotic potential as humans (incident hosts) have been found infected with different clinical pictures: fever and rash in Africa (Ashraf et al. 2015), neuroinvasive infections (Gaibani and Rossini 2017) in Europe (meningoencephalitis (Cavrini et al. 2009; Pecorari et al. 2009); idiopathic facial paralysis (Simonin et al. 2018)). However, these manifestations remain anecdotal and the asymptomatic form appears to predominate, as demonstrated by incidental findings during screening of asymptomatic blood donors and seroprevalence studies (Angeloni et al. 2023).

In Africa, USUV seems not to be pathogenic for local bird populations, either because it is not a naturally virulent virus, or because of immunity or genetic resistance of birds, which have been in contact with related flaviviruses for a long time (Bakonyi et al. 2004). In Europe, common blackbirds are particularly affected, and to a lesser extent some birds of prey (Giglia et al. 2021), such as the great grey owl (Clé et al. 2019). The lesion patterns of 160 common blackbirds in the Netherlands included hepatosplenomegaly (major symptom), coagulative necrosis, lymphoplasmacytic inflammation and vasculitis. Reported symptoms included non-specific ones (immobility, apathy and ruffled plumage) and neurological ones (depression, stiff neck, inability to fly and epileptic seizures) (Giglia et al. 2021).

In several European regions, epidemiological patterns increasingly suggest endemic profiles rather than repeated introductions from other endemic countries (Constant et al. 2022), without clear explanation of efficient overwintering and subsequent amplification (Clé et al. 2019). Moreover, co-circulation with WNV is frequent (Beck et al. 2013; Fros et al. 2015; Constant et al. 2020). These situations have led European countries such as Italy to institute annual surveillance (entomological, human and veterinary), rather than focusing on a specific season (Constant et al. 2022).

In France, the first virus detection (in August and September 2015) took place in the North-east (Haut-Rhin department) and Central-east of France (Rhône department), respectively in two and three dead common blackbirds collected through SAGIR, the French event-based surveillance network collecting and analysing dead free-ranging wildlife animals (see details in next paragraph). Phylogenetic analysis helped demonstrating proximity between isolates i) from Haut-Rhin and Germany and ii) from Rhône and Spain. In addition, isolates identified in the two French departments belonged to different USUV lineages (Europe 3 in Haut-Rhin, Africa 2 in Rhône). Those results suggested a different source of introduction for the two French departments (Lecollinet et al. 2016). Since 2015, outbreaks in French avifauna have been recorded yearly in 2016, 2017 and 2018, with new departments involved (Johnson et al. 2018). In 2018, the Usutu virus circulated earlier and more extensively than it had in preceding years (Épidémiosurveillance Santé animale (ESA) 2018). In Camargue (South of France), which is located at the crossroad of several bird migration routes (Vittecoq et al. 2013)) two lineages (Africa 2 and Africa 3) were found in 2016 among female *Cx. pipiens*, suggesting two independent introductions of the virus: i) one from West to East and probably explained by migratory birds following the East Atlantic flyways (Africa 2 lineage) and ii) one explained by migratory birds following a central Mediterranean flyway and spreading after to Northern Europe, or originating from Germany (Africa 3) (Eiden et al. 2018). The USUV detection, in the southern region of Camargue, of same lineage, in same region and for consecutive years (2015, 2018 and 2020) in mosquitoes highly suggests endemicity (Constant et al. 2022).

Until now, studies on USUV in France focused on emergences and genetic characterisation to reconstruct introduction paths (Vittecoq et al. 2013; Lecollinet et al. 2016; Eiden et al. 2018; Constant et al. 2022), but not on spatial, temporal, spatio-temporal and populational aspects of the virus diffusion within the country. The aim of our study was twofold: i) from an epidemiological perspective, we first analysed the USUV diffusion among related bird populations to understand the way the disease had spread throughout mainland France and thus to define an epidemiologic pattern of this 2018 bird outbreak and its relationships with environmental variables (in particular wetlands, supposed to be a proxy for the density of mosquitoes); ii) from an ecological perspective, we tried to understand how this diffusion could have impacted common blackbird populations.

For the epidemiological perspective (both infectious and spatial approach), we used data from the SAGIR network which is dedicated to epidemiological surveillance, through the collection of dead or moribund birds and mammals (opportunistic sampling method). This participatory surveillance program aims at detecting as early as possible abnormal mortality or morbidity signals (Millot et al. 2017; Decors et al. 2022).

Moreover, in 2018, abnormal mortalities due to USUV have also been observed on captive birds in zoos and were reported in the database of the French Association of Zoo Veterinarians (AFVPZ). The French Agency for Food, Environmental and Occupational Health & Safety (Anses) combined SAGIR and AFVPZ data. This dataset offered an opportunity to study the epidemiology of USUV (date, location, and size of clusters), by identifying the timing and geography of reporting of dead birds positive to USUV in France.

We expected that in clusters with high density of dead birds identified from this dataset, the high mortality would have an impact on the population dynamics of wild birds. Common blackbirds, in particular, are of interest as this species is the most affected species by USUV in France. The impact of USUV on common blackbird populations can be assessed thanks to the data collected by another French observation network named “REZOP” (Réseau Oiseaux de Passage, passing birds network), which is dedicated to the evaluation of bird population dynamics and focuses on non-captive wild Alaudidae, Columbidae, Turdidae, Phasianidae, Corvidae and Sturnidae. In this program, birds belonging to the focus species are counted every year, through a systematic sampling method and collected data are statistically analysed to define population trends (Villers et al. 2021).

We hypothesised that USUV circulation in 2018 resulted in decreased common blackbird populations in the years following the 2018 epizootics, based on a previous study on blackbirds dead following USUV infection in Germany between 2011 and 2015 (Lühken et al. 2017). Although the approach of this German study was slightly different, as we will discuss in the dedicated part of this paper, we also expected that USUV circulation in birds would be associated with some environmental variables and would have a negative impact on common blackbird population trends.

## Materials and methods

### Study design

This study employed an observational ecological design, using data from existing surveillance networks to assess the spatial and temporal distribution of USUV infections among bird populations in France. Data were collected opportunistically from mortality events reported through the SAGIR and AFVPZ surveillance networks, encompassing wild and captive birds. The study design allowed for the examination of environmental and ecological variables influencing USUV distribution, such as proximity to wetlands and human population density, which serve as proxies for mosquito habitats and bird sampling pressures respectively. We also identified spatial patterns (i.e., clusters) and temporal trends of USUV outbreaks at the population level. We used REZOP data to assess the correlation between estimated USUV infection levels and observed trends in blackbird populations across various geographic areas.

### Data collection and surveillance networks

The birds in our study were collected through the SAGIR network and zoological gardens across mainland France (Fig. 1A) from July 15^th^ to August 31^st^, 2018 (after this latter date, a message was sent to the local coordinators of the SAGIR network, asking them to limit data collection to only one sample per department, thus avoiding the additional cost of analyses linked to the increased epizootic outbreak; data collected by the network were thus defined at the department level from September onwards). Before August 31^st^, the most precise geographical information for each reported case was the municipality (a municipality being the smallest French administrative subdivision, corresponding to town and village areas, with a median area of 9 km²).

**Fig. 1.**
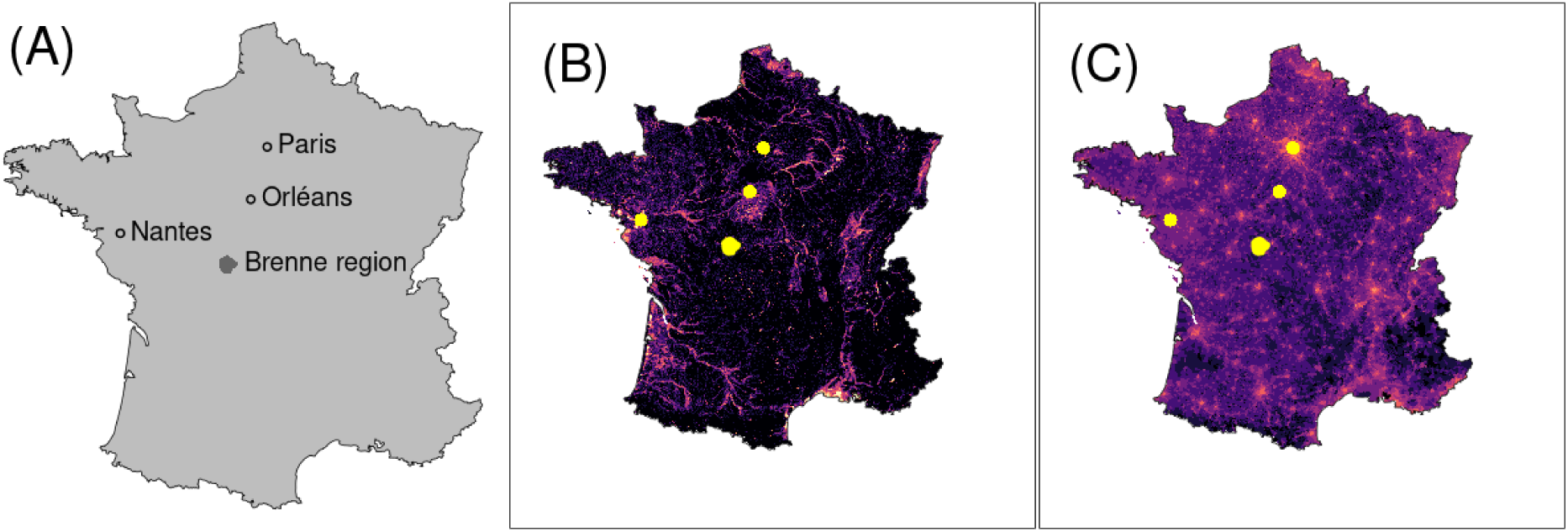
Study area (grey) (**A**) with regions and towns mentioned in the study, wetlands (**B**) and human population densities (**C**) (**B**: from black to pink: from the least likely to the most likely wetlands; **C**: from black to pink: from the least to the most densely populated areas; **B** and **C**: yellow surrounded by a yellow line: main towns and regions mentioned in the study)

Data were provided i) mainly by the event-based surveillance network SAGIR and ii) by the AFVPZ surveillance of dead animals in zoos (with systematic necropsy and analysis of additional samples depending on observed lesions and ante-mortem clinical picture).

SAGIR is based on a collaboration between the French Office of Biodiversity (OFB), who administrates the network, hunters, local and national federations of hunters and the French Association of directors and executives of public veterinary analysis laboratories (Decors et al. 2022). In each *department* (a department being a French administrative district covering in average 5,700 km² [SD = 2,500 km^2^]), two departmental technical contacts (one belonging to the departmental service of OFB, the other belonging to the departmental federation of hunters) (Decors et al. 2022) coordinate a network of volunteers (professionals, as well as hunters, naturalists and farmers) who can report abnormal events (mortality or morbidity) that they accidentally find in the field. The reported events can also include dead animals of local rescue centres on rare occasions. The sampling of these events is therefore opportunistic. For traceability purposes, a standardised individual digital form is filled for each mortality event collected, reporting epidemiological, agricultural and ecological circumstances surrounding the discovery of carcasses. When available, clinical signs are also provided (Millot et al. 2017), deduced from observation of the carcass and signs left in the environment. Sometimes animals are seen alive before they die (e.g. 12% of common blackbirds in the database from 2014 to 2023, regardless of cause of death) (A. Decors, personal communication).

### Laboratory procedures

In SAGIR, carcasses (fresh, chilled or frozen) accompanied by their form are addressed to the local administrative laboratory of veterinarian analyses where necropsies are realized. Following a gross pathologic examination, relevant aetiology tests are conducted: parasitology, bacteriology, virology, mycology, toxicology and/or histology (Millot et al. 2017).

In AFVPZ network, dead zoo animals are systematically necropsied within a maximum of 20 hours (usually less than six hours after death). The zoo veterinarian performs the necropsy and takes biological samples when lesions are observed, for analysis (bacteriological, mycological, parasitological, or virological if relevant). The samples are sent to the nearest departmental veterinary laboratory or, in some cases, to a laboratory specialised in the species concerned. Histological analysis is also carried out almost systematically.

Birds collected from both networks were also tested by the National Reference Laboratory (NRL, Anses), if i) their ante-mortem clinical picture was suggestive of an USUV infection (e.g. neurological symptoms) and/or ii) the epidemiologic context was consistent with a USUV suspicion (close mortality among sensitive species, and especially great grey owls) and/or iii) lesions at post-mortem examination were suggestive of an USUV infection (liver, spleen, cerebral necrosis). Spleen, liver, brain and lung (sometimes kidney and heart) samples were then sent to the NRL and tested for USUV and WNV through Real-Time Polymerase Chain Reaction (hereafter RT-PCR). Tissues samples were stored at −80°C before RT-PCR analyses and were grinded in Dulbecco modified Eagle’s minimal essential medium (DMEM) (Lecollinet et al. 2016) with ceramic beads (MP Biomedicals, Illkirch, France) and FastPrep ribolyzer in BSL3 facilities. Automated total RNA extraction, amplification and detection of USUV genome were carried out as described in (Moutailler et al. 2019).

### Epidemiological data analysis

In our study, we focused only on birds reported USUV infected, and we discarded other reported birds. We found it easier to analyse the space and time structure with point patterns (availability of a large variety of methods to deal with such data) so that we transformed our dataset into a point pattern. Thus, for each reported bird, we randomly attributed a precise point location by randomly selecting a point within the municipality where it was collected. Hereafter, the terms ‘points’ or ‘cases’ will refer to these birds. All USUV-infected birds were considered, whatever their species or origin (captive versus non-captive).

We analysed the temporal, spatial and spatio-temporal distribution of reported cases, as well as the relationship with spatial clusters of cases with wetlands and with human density (Fig. 1B and C). Wetlands and human density were supposed to be a proxy for the density of *Culex pipiens* mosquitoes (Becker et al. 2010; Haba and McBride 2022). Human density was also supposed to be a proxy for common blackbirds sampling pressure, as those birds are highly commensal with humans and easy to detect in gardens. A map of potential wetlands was derived from a raster map of continental France built by INRAE (Orléans) (http://geowww.agrocampus-ouest.fr/web/?p=1538): the INRAE map estimates the probability that each 1km x 1km pixel of the map harbours a wetland and then discretize this probability in three classes (0 = no wetlands, 1 = rather strong probability, 2 = strong probability, or 3 = very strong probability). This map also identifies lakes and foreshores in France. We downloaded this map, and redefined pixels with lakes and foreshores as “confirmed wetlands” (new class 4). Then, we smoothed this map using a sliding window, by calculating for each pixel the sum of this ordered factor (0 to 4) for the focus pixel as well as the pixels immediately on the left, the right, the top and the bottom of the focus pixel (leading to a score comprised between 0 and 20 giving a smoothed probability of wetland in a given place; see Fig. 1B).

A map of human density was calculated by finding the municipality in which the centre of each 1km x 1km pixel was located and attributing the 2005 human density to it (number of inhabitants divided by municipality area; see Fig. 1C). We then log-transformed this map (transformation: *x → log(x+1)* where *x* was the number of inhabitants per km²). The 2005 human density was provided by Institut Geographique National (France) (IGN) (https://geoservices.ign.fr).

We then carried out a statistical analysis of the SAGIR and AFVPZ data aiming at identifying spatial and spatio-temporal patterns. All our analyses were carried out with the R software (R Core Team 2022). We have programmed an R package named usutuFrance, available at https://github.com/ClementCalenge/usutuFrance, containing all the code and data used to fit the model. It can be installed in R with the package devtools, using the function `devtools::install_github(“ClementCalenge/usutuFrance”, ref=“main”)`. This package includes a vignette describing how the user can reproduce the calculations carried out in this paper (vignette available with the command vignette(“usutuFrance”) once the package has been installed and contains supplementary analyses that helped to understand the structure of our data. This vignette serves as the supplementary material of our paper.

#### Spatial analysis

We first used a Ripley’s K function to identify the scales at which clusters of points could be identified (Diggle 2013). This function allows the characterisation of second order point pattern properties: it is proportional to the mean number of other cases expected in a *x* km radius around a typical case of this point pattern (so that the larger it is, and the more there are cases in the neighbourhood of a case in average – in other words, clusters can be observed at this scale). The K function was computed for all distances *t* comprised between 0 and 250 km on the observed point pattern 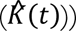; the theoretical value of this function was also calculated for the same range of distances under the hypothesis of complete spatial randomness (CSR) (*K*_*CSR*_(*t*)). Thus, under the hypothesis that the cases are randomly distributed in space, the difference 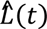 between 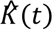 and *K*_*CSR*_(*t*) should be equal to 0 (Baddeley et al. 2014), so that positive deviations of 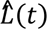 from zero reflect clustering in the data (and conversely, negative values reflect repulsion mechanisms). We also built confidence envelopes around simulated *L*_*r*_(*t*) functions expected under CSR: we repeated n=100 simulations of the CSR (by randomly distributing *N* cases in France, where *N* is the observed number of cases in our study), and for each simulation *r*, we estimated the function *L*_*r*_(*t*) for all values of *t* from 1 km to 250 km. We plotted the observed value 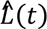 and the distribution of values expected under the CSR hypothesis on the same graph to compare them (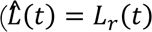: complete random reporting of cases; 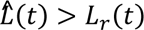: more points within a *t* radius around a typical case than under CSR hypothesis; 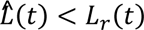: less points within a *t* radius around a typical case than under CSR hypothesis i.e. avoidance process).

To identify areas of high density of cases in the point pattern, we carried out a kernel smoothing of the point pattern, using a smoothing parameter of 125 km (Wand and Jones 1995), to visualize them on a map. The results (see Results section) suggested that the point pattern could be adequately described by a Thomas process (Diggle 2013), i.e. a process in which cases are clustered in the following way (Baddeley et al. 2014): i) a random number of “parents” is generated from a Poisson distribution parameterized by a parameter κ controlling the density of ‘parent’ cases per km². These parents are randomly placed in the study area. ii) For each “parent”, a random number of “children” (corresponding to the cases in our study) is generated from a Poisson distribution parameterized by a parameter λ controlling the mean number of “children” per “parent”, and iii) for each “parent”, the children locations are randomly distributed around the parent’s location.

More precisely, the locations of the children are supposed to be located at a distance from the parent randomly drawn from a semi-normal distribution with a standard deviation σ, with an angle from west direction drawn from a uniform distribution bounded by π and -π. We fitted this process to our dataset using the method of minimum contrasts based on the function *K*. Estimated parameters of the fitted model helped defining the point pattern typical structure and the product of λ and κ provided the cases density per km². Note that our spatial analysis revealed a clustering of the dataset at very small scale (see results), possibly due to the tendency of observers to look for other cases in a municipality where a first case was identified. Consequently, and to improve model fit, we decided to thin the point pattern prior to the fit to remove this artefactual structure, keeping only the first bird found when two birds were collected at a distance lower than 5 km (see Supplementary material for more details).

#### Environmental determinants

Then, we used this fitted model to test, using a randomization approach, whether the mean value of the two environmental variables (wetlands and population density) were greater in the places where the dead birds infected by USUV were found than expected by chance (Manly 1991): i) we first calculated the means of the density of wetlands and the mean of the log of human population density within the observed points pattern, ii) we simulated the fitted Thomas process 999 times over France, iii) we calculated for each simulation the mean of the density of wetlands and the mean of the log of human population density (simulated distribution), and iv) we compared the observed values (means) with the simulated distribution (means and standard errors) and determined p-values with the help of a bivariate randomization test.

#### Temporal analysis

To define the temporal evolution of the number of cases throughout the study period, we computed the mean number of cases per day during this period and smoothed the curve using moving average approach (Diggle 1990): for each *d* day of the study period, we calculated the mean number of cases per day, over a week centred on *d*. We plotted the obtained values on a graph to visualize temporal fluctuations.

#### Spatio-temporal analysis

We computed a space-time K function 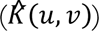 (Gabriel et al. 2013) giving the expected number of cases within a distance of *u* km around a randomly sampled case of the observed point pattern (for all values of *u* comprised between 0 and 250 km), and which occurred at most *v* days after this case (for all values of *v* comprised between 0 and 10 days). To test whether the spatio-temporal patterns identified by this function could have been obtained by chance, we also used a randomization approach. More precisely, for each one of the 1,000 simulations of this approach, we randomly permuted the dates associated to the reported birds (i.e. the cases order was randomised), thus conserving the spatial and temporal point pattern structures but randomising the association between space and time. For each simulation, we calculated the related *K*(*u*, *v*) on the randomized dataset. We then compared, for each distance *u* and each duration *v*, the observed value of 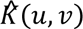 to the distribution of simulated values *K*(*u*, *v*). Probabilities that a simulation could lead to 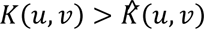 were plotted, with duration on the x-axis, distances on the y-axis (probabilities near zero indicating clustering) and a contour limit identifying the set of distance/duration pairs for which the proportion of simulation with 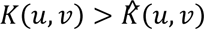 was lower than 5%. This allowed to identify the space-time clusters in our dataset.

### Impact on common blackbird population densities

#### Bird survey and data collection

The abundance of common blackbirds was monitored in France over the 1996-2022 using a large-scale survey designed to assess population trends of common game species (*ACT Survey* (Boutin et al. 2001, 2003)). The spatial coverage of this survey is based on 1,070 grid cells measuring 28 x 20 km (1/50 000e IGN map; see Supplementary material). Each cell includes a sampling route (∼4 km in length) randomly selected outside highly urbanized areas, with five points counts spaced by ∼1km. Each year, these points are surveyed twice during the breeding season by hundreds of observers (affiliated to OFB or departmental federation of hunters): from April 1^st^ to 30^th^ and from May 15^th^ to June 15^th^. The census protocol adopts the point count methodology: each point is surveyed for ten minutes in early morning. During this period, observers record the number of different singing males within a 500 m radius circle around the point (see Villers et al. 2021 for more details). Over the 1996-2022 period, each route was surveyed on average for 24.6 years (SD +/- 3.8 years; range: 1-27) and each year, the number of surveyed route was on average 977.1 routes (SD +/- 69.5; range 750-1038).

#### Population trends

For a given route and survey date, the number of singing birds was summed across the five points. We ensured that for each observation, all the following conditions were met: i) time of the count was available, ii) the start time of the survey was between −60 mn and +180 mn relative to sunrise time and iii) the date of the observation was posterior to the 31^st^ of March and anterior to the 30^th^ of June. We used these date and time intervals as presumed windows that minimize biases related to the detection rate of singing males. Our final dataset included 1,070 different routes (see Supplementary material), totalizing 50,454 counts over the study period.

We used the framework of Generalized Additive Mixed Models (GAMM) implemented in the R package mgcv (Wood 2006) to calculate annual population indices and underlying population trends. The number of singing males per route and per date (*Counts*) was modelled as a function of a smoothed function (using thin plate regression splines) of the counting *Date* (expressed as a Julian date, i.e. the number of days elapsed since the beginning of the year), a smoothed function of *Time-of-day* (expressed in minutes relative to local sunrise thanks to the R package suncalc (Thieurmel and Elmarhraoui 2022), and a random effect associated to *Route ID* (a 1,070-levels factor). We used *Date* and *Time-of-day* as smooth terms to capture some components of the detection process (Knape 2016), including plausible change in singing rates in relation to breeding phenology and/or singing behaviour during the morning. In order to straightforwardly compare trends in the abundance index according to different levels of USUV incidence, we also created a factor *Usutu.Area*, indicative of the severity of the 2018 USUV episode in space. For each point of the *ACT Survey* we extracted the value of the density of USUV cases estimated by the kernel smoothing carried out previously (cf. supra, section USUV diffusion process, Spatial analysis). These values were averaged for each *route* (mean value of the five sampling points), and converted into a 3-levels factor, according to the 50^th^ and 75^th^ percentiles of the distribution of USUV cases density: *routes* displaying kernel values below the median were coded as *low*, over the 75^th^ percentile as *high,* and *medium* otherwise. In order to model the temporal variation of the abundance index in each area, we included the *Usutu.Area* as a fixed effect, a smooth function of years (therefore considered as continuous covariate, *Year,* with a thin plate regression spline), and an interaction term allowing to fit a temporal trend to each level of the *Usutu.Area* factor (with independent smoothing parameters (Pedersen et al. 2019)). This interaction allowed to identify different population trends corresponding to different levels of USUV cases density. Finally, we also included in the model a random effect associated to the year considered as a 27-levels factor (*Year.as.Factor*). This statistical approach allowed separating short-term fluctuations (from one year to the next) from longer-term (over a decade) population changes, through the inclusion of temporal random effects in simulation of smooth trends (Knape 2016). The number of degrees of freedom of the basis function of the smoother of the year effect was set equal to k=10.

Models were fitted with a Tweedie distribution. Tweedie distributions, a special cases of exponential dispersion models, are probability distributions encompassing the continuous (normal, gamma and inverse Gaussian), discrete (Poisson) and compound (Poisson-gamma) distributions. In the R package mgcv, the value of the power parameter *p* of the Tweedie distributions (with 1 < *p* < 2) can be estimated simultaneously with the estimation of other parameters, offering flexibility in the type of response that can be modelled.

To compute the trends and their associated confidence intervals from the fitted model, we adopted a posterior simulation *sensu* (Wood 2017) (chapter 6.10, p. 293), in order to obtain the distribution of predicted values of abundance for each level of the *Usutu.Area* factor. A similar approach is implemented in the R package poptrend (Knape 2016), but it does not offer the possibility to compute trends for specific models, e.g. aiming at comparing trends for different levels of a grouping variable. The general idea is to combine prediction matrices and simulation (replicate) from the statistical distribution of the parameters *β* of the model, in order to obtain distribution estimates (i.e. including uncertainty of parameter estimate) of any quantity of interest, in our case the relative abundance index of common blackbirds for an average route and year, hereafter referred to as the variable *Index*. Details of the computations can be found in the supplementary material.

Trends were expressed as

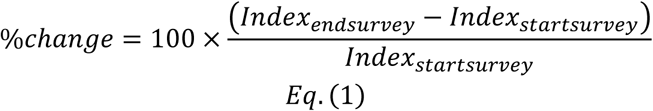

To estimate population trends accounting for the variability of the abundance index, we computed, for each simulated replicate, the mean abundance index for the four years before (2015-2018) and after (2019-2022) the USUV episode of the 2018 summer. We then plotted for each level of the *Usutu.Area* factor the value of the abundance index for the two periods (2015-2018 and 2019-2022) and compared the population trends for each area thanks to Eq. (1).

## Results

### USUV diffusion process

#### Epidemiological surveillance network and provided data

The dataset (60 % SAGIR, 34% AFVPZ and 6% rescue centres) included 60 birds tested for USUV of which 50 were detected infected in 29 French departments between July 15^th^ and August 31^st^, 2018. Wild common blackbirds and captive great grey owls were particularly affected even if several other species were concerned (Tab. 1).

**Tab. 1.**
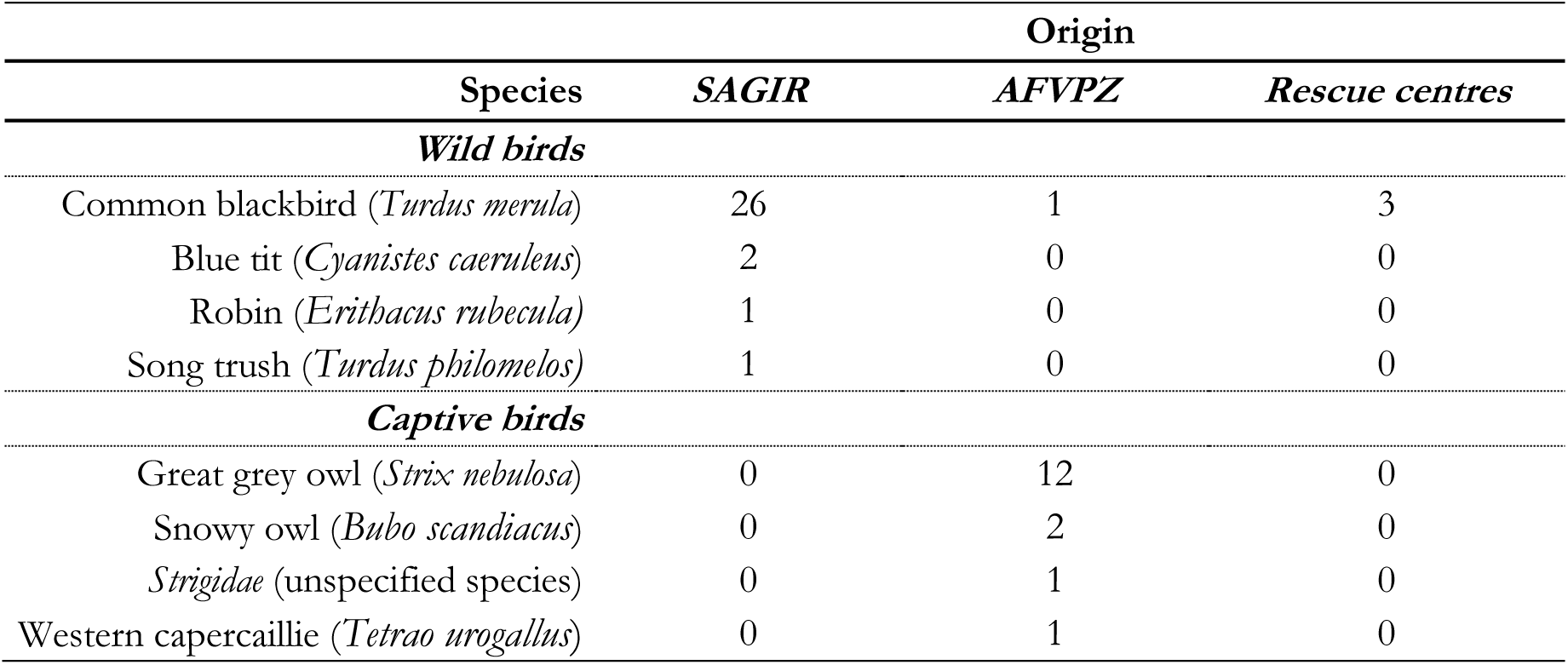
Distribution of birds collected and detected infected by USUV within the two epidemiological surveillance networks of the study between July 15th and August 31st, 2018 (data from rescue centres were stored in the SAGIR database).

#### Spatial analysis

The comparison of observed *L*^^^(*t*) and the distribution of simulated *L*(*t*) (n=100) showed that infected birds were not randomly distributed and were concentrated in specific geographic areas: it allowed to identify a clustering of the point pattern at 5 km, 125 km and 200 km (Fig. 2A).

**Fig. 2.**
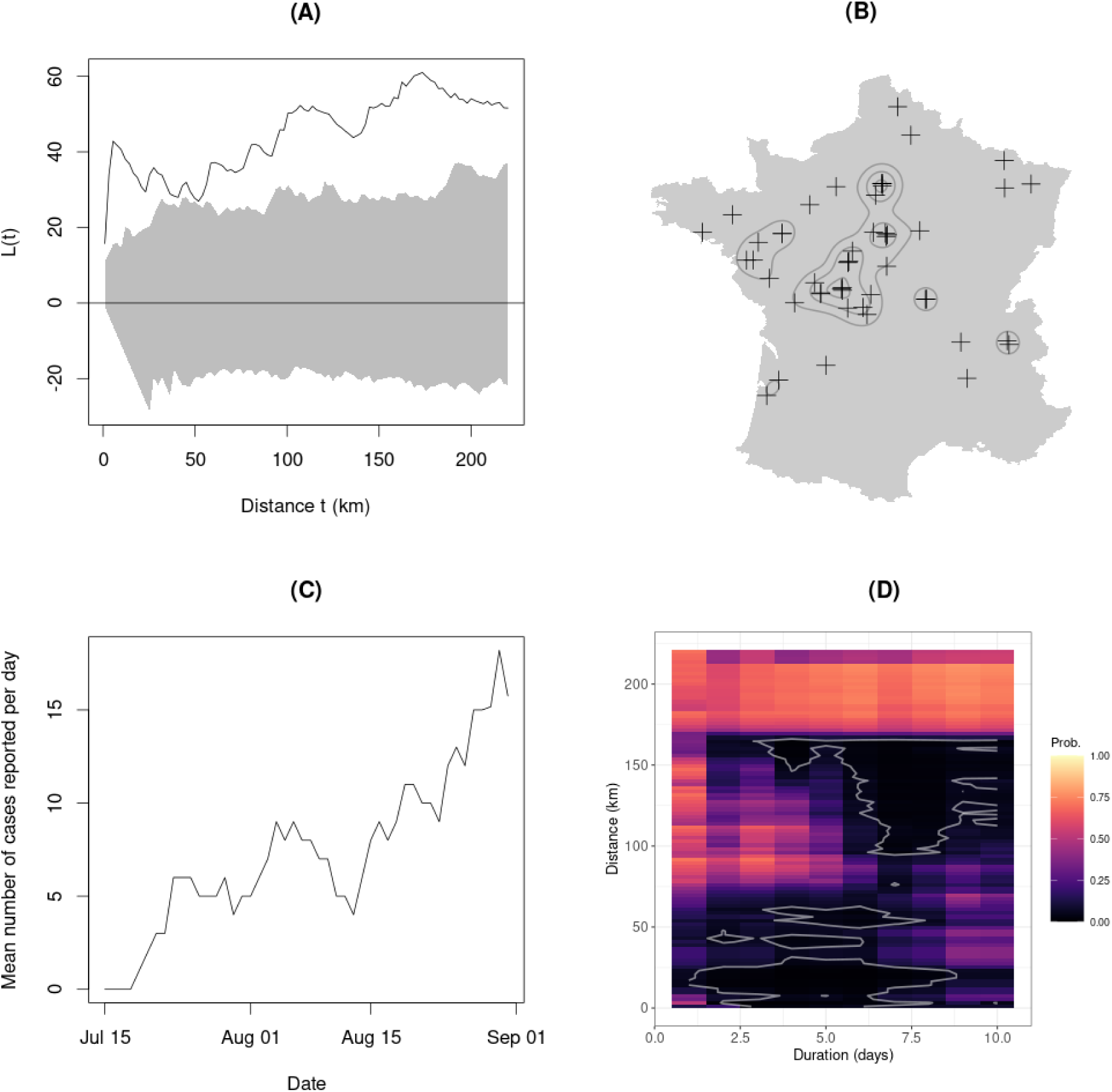
Spatial (**A**, **B**), temporal (**C**) and spatio-temporal (**D**) characteristics of the point pattern of birds detected infected to USUV during the study period (**A**: Comparison between 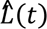 (observed, grey line) and *L*_*r*_(*t*) (envelope of simulated point pattern under CSR hypothesis, grey area); **B**: Kernel smoothing of the point pattern (grey lines) and birds locations (black crosses); **C**: Mean number of cases (mobile average approach) reported per day and depending on date; **D**: Probabilities that spatio-temporal K function *K*(*u*, *v*) simulated under the hypothesis of independence between spatial and temporal patterns, was higher than observed spatio-temporal K function 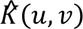, depending on duration and distance (white line: contour limit of the set of distance/duration pairs for which the proportion of simulation with 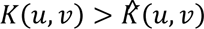 was lower than 5%)

The kernel smoothing carried out with a smoothing parameter equal to 125 km showed a high density of reported cases in the Brenne area (largest cluster located at the southwest of Paris on Fig. 2B – Paris being the northernmost cluster delimited by contour lines on the figure (see Fig. 1A for the names of towns and regions). Other high cases density areas could be identified, particularly in Paris, towards Orléans (south of Paris) and Nantes (west of France; Fig. 1A, Fig. 2B). The distance between Paris and Orléans high USUV cases density areas is around 200-250 km, as well as between those of Brenne and Nantes. This last result was consistent with the third peak observed in Fig. 2A (clustering at 200 km).

The fitting of the Thomas process model helped determining the density of clusters (“parents”) per 10,000 km^2^ (κ parameter: 0.17) and the mean cluster size (λ parameter: 4.36, mean number of “children” per “parent”). Thus, the mean density of detected cases was estimated to be 0.74 cases per 10,000 km^2^. The standard deviation of the distribution of distances between the cases and the centroid of the cluster (the parent case) was of 77 km. This last result was consistent with the K function’s second peak, which indicated the presence of clusters at a scale of about 130 km. Indeed, this standard deviation measures the typical distance between cases and the centroid of the cluster, while the K function is based on the distance between two cases. By simulation, we checked that the mean distance between points sampled in a bivariate Gaussian distribution with a standard deviation of 77 km was close to 130 km (see Supplementary material).

#### Environmental determinants

The observed mean of wetland probability score in USUV places was equal to 4.81, while the one expected under the absence of effect of the environmental variables (for n=999 simulations of the fitted Thomas process) was of 2.69 (SD=0.65). The difference was significant (p-value=0.007), indicating that USUV cases were more often reported in or close to wetland than expected by chance. This mean wetland probability score in USUV places corresponded to the top 19.3% most likely wetland areas. Similarly, the observed mean of human log-density population in USUV places was of 5.02, while the one expected under the hypothesis of absence of effect of the environmental variables was of 3.50 (SD=0.25). The difference was also significant (p-value=0.001), indicating that places where dead birds positive to USUV were reported were more often located in places with high human density. This mean log-density population in USUV places corresponded to the top 10.5% densest areas in France.

#### Temporal analysis

There was an initial increase in the number of cases between July 15^th^ and 20^th^ 2018 (Fig. 2C). Then from July 20^th^ to August 15^th^ the number of cases reported per day stagnated at about one per day. From August 15^th^ onwards, there was a sharp increase, with an average of more than two cases reported per day towards the end of the month.

### Spatio-temporal analysis

The plot of probabilities that a simulation could lead to 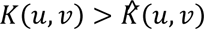 showed a strong clustering at small distances of about 10 to 20 km as soon as two or three days after a case, expanding to 50-60 km five days after a case. But we also observed a clustering on larger distances (150 km) starting as soon as two or three days after a case, which increased to reach a large range of distances after a week (90 to 150 km) (Fig. 2D). The large distances clustering was consistent with the first phase of the diffusion (between July 15^th^ and 20^th^) (Fig. 2C), with the emergence of the main clusters of Brenne, Paris, Orléans and Nantes (Fig. 2B). The small distance of clustering was consistent with the emergence of secondary clusters and the expansion of preexisting main clusters (Fig. 2B) during the third phase of the diffusion (from August 15^th^) (Fig. 2C).

### Impact on common blackbird population densities

The means and confidence intervals depicting cells of respectively low, medium and high USUV cases density have been reported in Tab. 2, the distribution of those cases in Fig. 3A and their spatial distribution in Fig. 3B.

**Fig. 3.**
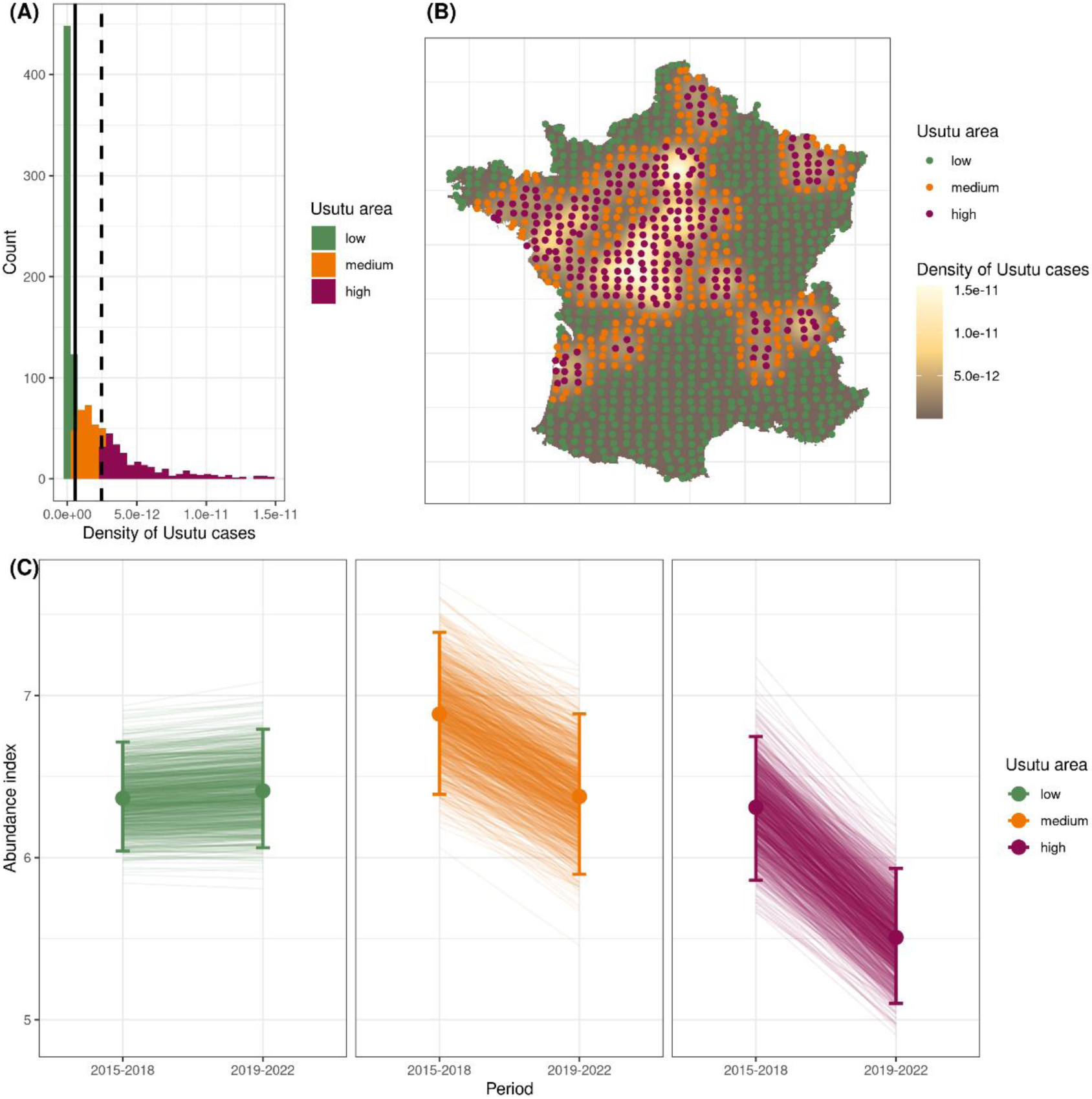
Common blackbird population trends in response to the USUV episode (**A**: Histogram of USUV cases density, from a kernel smoothing, for the routes where common blackbird were monitored (solid line: median of the distribution, dashed line: 75% percentile of the distribution); **B**: Maps of the kernel smoothing of USUV cases and locations of the centroids of REZOP road (pale green: low, orange: medium and dark purple: high densities of USUV cases as estimated by the kernel method); **C**: Predicted abundance index (mean values and their 95% confidence intervals) for the three classes of USUV cases density (pale green: low, orange: medium and dark purple: high densities) before (2015-2018) and after (2019-2022) the 2018 USUV outbreak. Each solid line represents changes in abundance value for a given simulation (1,000 simulated values were generated).

**Tab. 2.**
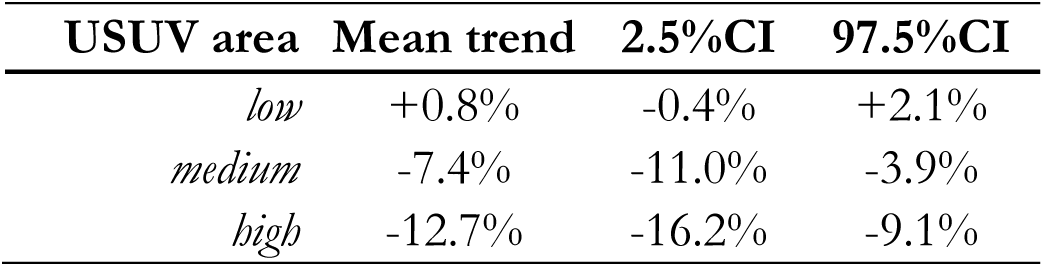
Trends computed between 2015-2018 and 2019-2022 for the three different levels of density of USUV common blackbird cases (CI: Confidence Interval).

| USUV area | Mean trend | 2.5%CI | 97.5%CI |
| --- | --- | --- | --- |
| *low* | +0.8% | -0.4% | +2.1% |
| *medium* | -7.4% | -11.0% | -3.9% |
| *high* | -12.7% | -16.2% | -9.1% |

The estimation of common blackbird population trends estimated through variations of the mean population indexes for 2015-2018 and 2019-2022 periods was not statistically different from 0 for the low USUV area but presented a significant decline for both medium and high USUV areas, being more pronounced for the latter than for the former (Tab. 2). Two main results were highlighted: i) a decreasing trend for common blackbird populations that have been *a priori* exposed to USUV since the 2018 USUV outbreak and ii) the greater the pressure of infection was, the greater the negative effect on common blackbird populations was (Fig. 3C).

## Discussion

Birds affected by USUV between July 15^th^ and August 31^st^, 2018, in France were mainly common blackbirds and to a lesser extent captive great grey owls. Cases were clustered at i) 5 km, ii) 125 km (which was the average distance between two typical cases which we highlighted with a Thomas process model) and iii) 200 km (which was the distance between main big clusters which appeared simultaneously at the beginning of the outbreak and expanded thereafter). Three periods of USUV diffusion have been featured: i) a first increase of USUV cases between July 15^th^ and 20^th^ (with clustering at large distances of about 150 km between two and three days after a case and between 90-150 km a week after a case, corresponding to the emergence of main clusters – Brenne, Paris, Orléans and Nantes), ii) a stagnation between July 20^th^ and August 15^th^, and iii) an important increase from August 15^th^ (with strong clustering at small distances of about 10-20 km, between two and three days after a case and expanding to 50-60 km five days after a case, corresponding to the emergence of secondary clusters and the expansion of preexisting main clusters). Detected case locations were statistically associated with wetlands and high human log-density population areas. Common blackbird population trends estimated suggested an impact of the USUV infection on them, with a decrease of the mean population index over 2015-2018 compared to 2019-2022. Areas of *a priori* greater USUV pressure presented more pronounced negative trends than the two other areas.

Species involved in the 2018 outbreak in the present study were those also found in other European countries such as Germany (Becker et al. 2012; Lühken et al. 2017). Several hypotheses might be put forward regarding the overrepresentation of common blackbirds: i) they are not trans-Saharan migratory species and as a consequence probably deprived of an immunity they could have acquired in Africa, ii) they might be more genetically susceptible than other birds species, iii) African strains introduced in Europe might have adapted to common blackbirds and become more virulent for this host (Bakonyi et al. 2004), iv) co-infections with *Plasmodium* might exist with a possible interplay of the two agents, as highlighted in the Netherlands (Agliani et al. 2023), v) a feeding preference has been identified for them (when they were sufficiently abundant) by *Cx. pipiens* (Rizzoli et al. 2015) and vi) this species is abundant, ubiquitous, human commensal (Lühken et al. 2017) and thus easy to detect by the SAGIR network.

Any pattern identified by the analysis of data collected by a participatory program relying on non-controlled sampling pressure can be caused by the opportunistic observation process (characterized by a very heterogenous sampling pressure) as well as by the ecological or epidemiological process. In particular, the data collection process differs across departments (differences in detectability, in probability to report a carcass to the network, in probability to collect this carcass by the local contact, mode of transport, mode of storage, etc.). Thus, when a pattern was identified in the data (e.g. a cluster of cases), we could not decide between (i) the fact that this pattern was caused by the observation process (e.g. higher local sampling pressure due to higher human density and thereby detectability) and (ii) a biological cause (e.g. local factors favouring the diffusion of USUV in this place). It would have been interesting to model the data collection process to draw firmer conclusions on the biological process. However, the lack of information about the data collection process within the SAGIR network has been a long-standing issue, hindering such modelling (as is often the case with participatory networks). Our approach was therefore exploratory, and our results could only help formulating hypotheses, and could not allow confirming them. Keeping in mind this limit, we must stress that our results pointed out that such a network and statistical analysis of admittedly imperfect data could however allow the understanding of diffusion pattern and epidemiological process. Indeed, the network was alerted to the possible presence of USUV cases in France and the level of attention given to surveillance was greater for this disease than for other diseases. We considered the possibility that any pattern identified in our data could be caused either by the observation process (e.g. the heterogenous sampling pressure) or by ecological/epidemiological processes.

The short distance clustering (at 5 km) identified by our analysis could be explained by local sampling effort. We could not exclude that observers would have been more proactive in detecting dead birds around first detections. However, this short distance of 5 km is also close to flight distances of *Cx. pipiens* estimated in a study carried out in Chicago, Illinois, on *Culex* female mosquitoes. Indeed the mean dispersal distance was 1.15 km, and 90% of individuals stayed within 3 km from their larval habitat (Hamer et al. 2014). Another explanation for the 5 km clusters we observed would be a USUV diffusion mainly due to mosquitoes. Sufficient amplification of the virus in its various hosts would also have allowed a large number of mosquitoes to be infected.

The occurrence of different clusters simultaneously at the beginning of this 2018 USUV outbreak, far apart from each other, is in favour of USUV endemicity throughout the country. The endemicity has already been proven in Southern (Constant et al. 2022) and North-Eastern France (Johnson et al. 2018), but the 2018 outbreak we studied and the diffusion pattern we showed raised the hypothesis of a national endemicity. Previous outbreaks, even of lesser magnitude, might have enabled certain lineages to settle. Indeed, in France, USUV Europe 3 strain has been identified in common blackbirds in two north-eastern departments between 2015 and 2017. USUV Africa 2 strain has been isolated in common blackbirds, mosquitoes and humans in three south-eastern departments and in Rhone between 2015 and 2016, while USUV Africa 3 strains have been identified in mosquitoes and birds in two central and four south-eastern French departments between 2015 and 2017 (Johnson et al. 2018, NRL data). Favourable conditions (e.g. sizes, naive immune status and/or turnover of bird populations, temperature, rainfall, human population movements) could then lead to new outbreaks thereafter. The early detection of this 2018 outbreak was made possible by the well-informed SAGIR network, which vigilance has been reinforced since the 2015 outbreak for USUV relevant species. Throughout Europe, endemic circulation and spread of USUV infections have been reported, e.g. Northern Italy, Hungary or Austria (Constant et al. 2022). The study of lineages involved within this 2018 outbreak could help validate endemicity.

The different phases of the 2018 USUV diffusion we observed remained difficult to explain: a first increase of cases, with clusters far from each other, a stagnation and then an important increase of cases, with secondary clusters and the enlargement of the first ones. If endemicity might explain the cases pattern of the first phase, the origins of the third phase remain unclear. Meteorological variables, as mean temperature (Hamer et al. 2014), could explain this result. Indeed, the duration of *Cx. pipiens* eggs development depends on temperature (hatching after only one day at 30°C, three days at 20°C, ten days at 10°C and incomplete embryonic development below 7°C (Becker et al. 2010)). Temperatures were particularly high in July and August 2018 in France (respectively +2.18°C and +1.1°C higher than the 1991-2020 baseline average (https://www.meteocontact.fr/climatologie/france/bilans-climatiques)). *Cx. pipiens* density might have increased in August due to these favourable conditions and promoted an increased possibility of contact between common blackbirds and the arthropod. Moreover, laboratory experiments have shown that higher temperatures resulted in higher infection rate of *Cx. pipiens* (Fros et al. 2015)). These conditions could have allowed an important development of *Cx. pipiens* and the third phase of USUV diffusion we could observe. Further studies collecting local relevant meteorological data would be useful to assess this hypothesis. Nevertheless, we could not exclude that the diffusion of information and knowledge of the presence of USUV in France would have also led to better sampling effort from August 15^th^.

In addition to the above-mentioned hypothesis related to the population dynamics of the main vector of the disease, other non-mutually exclusive hypotheses could explain the appearance of USUV clusters in late summer (after August 15^th^). This period coincides with the initiation of movements in common blackbirds, particularly juveniles that disperse outside their parents’ territories (Snow 1958), and to a lesser extent, with the first mentions of post-breeding migrants (Toulotte et al. 2022). Similar movements have been shown as the main drivers of seasonal epizootics of the avian influenza in mallards (van Dijk et al. 2014), and the same processes might be at play in the USUV episode reported here. USUV naïve birds such as juveniles, or individuals whose original populations have not yet been exposed to USUV, might be more prone to develop and spread the virus, easing the apparition of clusters.

In resident blackbirds, this period also overlaps with resource-demanding activities including breeding for some late breeders (Sauvage 2016), post-breeding (complete) moult in adults (Snow 1969; Morrison et al. 2015) and partial moult in juveniles (Cramp 1988). Previous works have shown that such activities may induce trade-offs with immunity in birds (e.g. Moreno et al. 1999; Sanz et al. 2004; Martin II 2005; Moreno-rueda 2010). Accordingly, late summer may coincide with a period of greater susceptibility to infections in blackbirds. Unfortunately, the lack of available data on age, sex, or moulting status, both for the recorded USUV cases, but also for individuals in the exposed populations, did not allow us to confirm these assumptions. This constitutes a relevant area of investigation for future studies and highlights the need for a deeper description of specimens found dead.

The fitted Thomas process was a simple model, with the assumption that all clusters were of the same size, that the mean number of cases per clusters did not vary from one cluster to another and that the density of “parents” generating secondary cases was uniform across the study area. However, we could not highlight any lack of fit of this model to our dataset, and by simulating this process we were able to simulate a realistic point process and thereby test the effect of environmental variables on the occurrence of USUV. This test showed that the density of USUV cases varied as a function of wetlands and human density, two variables that we supposed to reflect mainly mosquito abundance.

USUV is assumed to be mainly transmitted among birds by mosquitoes during their blood meal (Lühken et al. 2017), but a transmission from bird to bird could not be excluded in the diffusion process and the endemicity. Oral route transmission was suggested by damage of cells of the gastrointestinal tract observed as part of necropsies of infected common blackbirds and possible vertical transmission could not be excluded with antigen detection in the genital system (Giglia et al. 2021). An oral transmission could explain distances between cases higher than maximum flight distances of mosquitoes, while vertical transmission could be involved in endemicity mechanisms and perhaps even in the resurgence of cases after the reproduction period (from August 15^th^).

The association between wetlands and USUV was consistent with the life cycle of USUV, with mosquitoes as vectors and wetlands known as suitable habitat for *Cx. pipiens* (Becker et al. 2010; Haba and McBride 2022). The association with high human population densities could be more surprising at first sight. In an event-based network, the number of observers could partly explain such a result. Indeed, the greater the number of people able to collect birds affected by the disease would be, the greater the potential for the disease to be detected in densely populated areas would be. In addition, common blackbirds are familiar to urbanised areas (Partecke and Gwinner 2007), and their size and colour (especially in males) are striking (Lühken et al. 2017): collecting USUV dead common blackbirds would be easier in such areas. We therefore could not exclude that the association between human high-density areas and USUV places could be explained by a sampling bias, although this hypothesis is difficult to confirm with our opportunistic data. Actually, as for wetlands, a selection for high human density areas by mosquitoes might be a part of the explanation for this association with the USUV cases. Indeed, *Cx. pipiens* can also find favourable breeding grounds in urban (Haba and McBride 2022) or peri-urban (Vogels et al. 2016) environments (e.g. rainwater collection containers, ponds in gardens, flooded cellars or construction sites (Becker et al. 2010)).

In our study, data from the SAGIR surveillance network were used to determine USUV pressure areas on common blackbird population sizes, which have been estimated using GAMM in each class of USUV pressure. The combination of data from these two networks helped assessing the impact of USUV on common blackbird populations. The negative impact of USUV on common blackbird populations that we highlighted is consistent with what has been showed in Germany, where a difference of about 15.7% has been highlighted between the means of populations indices in 2016 in common blackbird populations (lower population index compared to the baseline year) and statistically significant lower population index in the USUV-suitable areas compared with the USUV-unsuitable areas (Lühken et al. 2017). Even if in our study the cut in three classes of USUV pressure helped us to highlight gradual effects on common blackbird populations tendencies (the greater was the infection pressure, the greater was the negative effect on common blackbird populations), our results and the German one converge in the conclusion that common blackbird populations are affected by the USUV circulation. This result points out the impact of USUV circulation on the balance of ecosystems including common blackbirds and the services they provide (e.g. a role in seed dispersal) (Whelan et al. 2008).

The presence of USUV infection in zoological gardens also raised the question of conservation of foreign species. As previously said, SAGIR network is a precious tool for early detection. But in case of endemicity, control measures must also be considered (e.g. elimination of artificial container habitats, reduction of shade sources over aquatic habitats (Tuten 2011), use of biological larvicides such as *Bacillus thuringiensis* subsp. *israelensis* (Virgillito et al. 2022), indoor caging during periods of high density of mosquitoes, mosquito nets). For the first time in 2018, the AFVPZ and SAGIR networks were really coordinated to face the USUV outbreak. Data collected and related to both captive and non-captive wild birds allowed the description and the characterization of the outbreak within mainland France (time of onset and duration). Non-captive wild common blackbirds could be seen in this context as USUV sentinel for captive wild birds and would be of great importance for species with conservation issues. This role could also be extended to non-captive other sensitive species (e.g. capercaillie or eagle-owl), and more specifically Passeriformes, as the birds collected in rescue centres have highlighted.

Another issue of the diffusion of USUV in France in wild birds as common blackbirds, living close to human populations, and in zoological species is contact with humans, who are known as USUV incident hosts (the asymptomatic form being predominant and fever illness, or neurological symptoms as encephalitis and meningitis (Vilibic-Cavlek et al. 2014) being rare (Angeloni et al. 2023)). Zoological parks are particularly favourable to the circulation of mosquitoes which can feed on both humans and animals (Martínez-de la Puente et al. 2020). In addition, screening of asymptomatic blood donors and seroprevalence studies have also revealed incidental findings of the virus in asymptomatic human populations (Cadar et al. 2017; Bakonyi et al. 2017; Zaaijer et al. 2019; Angeloni et al. 2023). Their link to epizootic outbreaks has raised the question of the need to strengthen USUV detection in blood donors, particularly during such periods (Cadar et al. 2017). Long-term epidemiological studies would be necessary to provide a better understanding of viral dynamics in potential wild reservoirs (Vittecoq et al. 2013). The USUV antigen detection in feather (follicle shafts and bulbs) of infected birds raised the possibility for live bird testing (Giglia et al. 2021). Risk mapping of competent vectors for USUV (*Cx. pipiens* and *Ae. albopictus*) could also help developing appropriate prevention and control measures (Martinet et al. 2023).

In a context of climate change (Walther et al. 2002; Parmesan and Yohe 2003), the impact on arboviruses is complex to define (Franklinos et al. 2019), but the perspective of a shorter wintering period and a longer period of activity cannot be ruled out. Such changes would not be without effect on the distribution patterns of arboviruses such as USUV and recent research has shown the role of climate change in the increase of the risk associated with West Nile Virus circulation through Europe (Erazo et al. 2024). Moreover, the analysis of the origins and the within-France spatial distribution of common blackbirds have highlighted that i) they mainly came from Western and Central Europe and ii) were strongly segregated in autumn and winter in France (Lahournat et al. 2021). The coming seasonal disturbances in a context of climate change therefore raise the question of the impact of USUV in France on its epidemiology on a European scale. The contributions of networks such as SAGIR and REZOP are therefore more necessary than ever.

Following this analysis of surveillance data, one could question the potential impact of the reactivity of a network such as SAGIR, coordinated with the other networks mentioned above, and what strategy should be adopted in the event of a future outbreak. Indeed, we highlighted here the relevance of combining data collected by different networks primarily dedicated to different purposes (e.g., SAGIR and REZOP). For example, an automatization of alerts could enhance the reactivity of the entire system of networks. The results of spatial analysis could be quickly accessible and help the definition of areas of risk, in near real time, and where surveillance (and if possible, control) efforts could be concentrated. Such an automatization could also be interesting in a financial perspective: as reported above, the number of cases had to be limited to one per department after August 31^st^, because of too high analysis costs associated with the growing outbreak. This drawback could be circumvented with better reactivity and if necessary, control of the virus diffusion. However, additional data (i.e. additional funding to analyse more samples even during the peak of the outbreak) would make it possible to reconstruct the spread of the virus over time by following different lineages more closely.

## Acknowledgements

We would like to thank all the observers and collectors of the SAGIR and REZOP networks, the local services of the French Agency for Biodiversity (OFB), the hunters and their departmental and national federations, the departmental laboratories for analysis and the national reference laboratory. We also thank the rescue centres and the zoological parks of the AFVPZ. We finally warmly thank Lorette Hivert and Nicolas Toulet for the Epifaune database managing and Jean-Philippe Martinet (Université de Reims Champagne-Ardenne, Faculté de Pharmacie, UR7510 ESCAPE–USC ANSES PETARD, Reims, France) for his valuable help in discussing entomological aspects and his review of these points.

## Compliance with ethical standards

In their publications, the beneficiary and associated partners specify that the work was carried out under the terms of an order derogating from the strict protection of species.

## Author contributions

CC performed the statistical analysis on USUV data. AV performed the populational statistical analysis on common blackbird populations. AD, BQ, AL and SL supervised USUV data collection. AV and CE supervised REZOP data collection. CC and AD interpreted results on USUV analysis. AV, CE, CC and AD interpreted both USUV and populational results. MBZ executed and reviewed the R code related to USUV diffusion and drafted the original manuscript. CC, AV, CE, SL and AD critically revised the manuscript. All authors contributed to the manuscript revision and approved the present version.

## Data, Scripts, code, and supplementary information availability

All the data, script, code and supplementary information have been packaged in an R package named usutuFrance, available on Github at https://github.com/ClementCalenge/usutuFrance. We have also stored this package on Zenodo (https://doi.org/10.5281/zenodo.10992191; (Calenge et al. 2024)). The raw dataset used in this paper has also been stored as a text file on Zenodo (https://doi.org/10.5281/zenodo.10992555;(Bouchez-Zacria et al. 2024)).

