## Supplementary material for "Relevance of the synergy of surveillance and populational networks in understanding the Usutu virus outbreak within common blackbirds (*Turdus merula*) in Metropolitan France, 2018"

### Appendices to the article: Analysis of the Usutu episode of summer 2018 in birds in France.

### Introduction

This vignette is the supplementary material of the article of Bouchez-Zacria *et al.* (prep). The aim of this paper is to analyze the space-time dynamics of the Usutu infection in French bird populations during the summer 2018, and to relate it with two environmental variables, and with the resulting changes in population size of blackbirds. A companion package named `usutuFrance` contains the data and R code used for this paper, and is required to reproduce the calculations in this document. The present document is also available as a vignette of this package. To install this package, first install the package `devtools` and use the function `install_github` to install `usutuFrance`:

```
## If devtools is not yet installed, type
install.packages("devtools")

## Install the package usutuFrance
devtools::install_github("ClementCalenge/usutuFrance", ref="main")
```

*Remark*: on Windows, it is required to also install the Rtools (`https://cran.r-project.org/bin/windows/Rtools/`) on your computer to have a working `devtools` package (see `https://www.r-project.org/nosvn/pandoc/devtools.html`).

Throughout this vignette, we suppose that the reader is familiar with the model and simulations developed in the main paper. We describe the R code used for this analysis, as well as additional checks that helped to better understand the dataset and underlying infection process.

### Appendix A Analysis of the SAGIR dataset

#### A.1 The datasets

In this appendix, we provide the R code used to analyse the infection process using the data collected by the SAGIR network. We first load the package:

```
library(usutuFrance)
```

We then load the data collected by the SAGIR network:

```
data(usutu)
str(usutu)

## tibble [60 x 5] (S3: tbl_df/tbl/data.frame)
##  $ species: chr [1:60] "Turdus merula" "Strix nebulosa" "Turdus merula" "Turdus merula" ...
##  $ date   : Date[1:60], format: "2018-07-22" ...
##  $ x      : num [1:60] 507007 599352 520995 579927 531686 ...
##  $ y      : num [1:60] 2188525 2428094 2144710 2317124 2274347 ...
##  $ result : chr [1:60] "Positive" "Positive" "Positive" "Positive" ...
```

This data.frame contains, for each one of the 60 dead birds collected by the network and tested for USUV:

- The latin name of the bird species (13 bird species);
- The date of collection;
- The x and y coordinates (Lambert II projection) of the bird;
- The result of the test (either "Positive" or "Negative").

In the paper, we focused only on the birds with a positive test result. Birds with a negative test result do not bring any useful information in this context: these are not dead birds randomly sampled over France, but rather birds for which local observers had strong suspicion that their death was caused by USUV:

```
usutup <- usutu[usutu$result=="Positive",]
```

We can show a map of these bird locations over a map of France. We load the dataset `sfFrance`, which is an object of class `sfc` containing the coordinates of the contour of mainland France; we then plot the locations on this map:

```
## Required libraries
library(sf)
library(ggplot2)
library(ggspatial)

data(sfFrance)

theme_set(theme_bw())
ggplot()+
    geom_sf(data=sfFrance)+
    geom_point(aes(x=x,y=y),data=usutup,size=3)+
    theme(panel.grid.major = element_line(color = gray(0.5), linetype = "dashed",
                                          linewidth = 0.5),
```

```
        panel.background = element_rect(fill = "aliceblue"))+
    theme(axis.line=element_blank(),
      axis.text.x=element_blank(),
      axis.text.y=element_blank(),
      axis.ticks=element_blank(),
      axis.title.x=element_blank(),
      axis.title.y=element_blank(),
      legend.position="none")+
    annotation_scale(location = "bl", width_hint = 0.5, plot_unit="m")
```

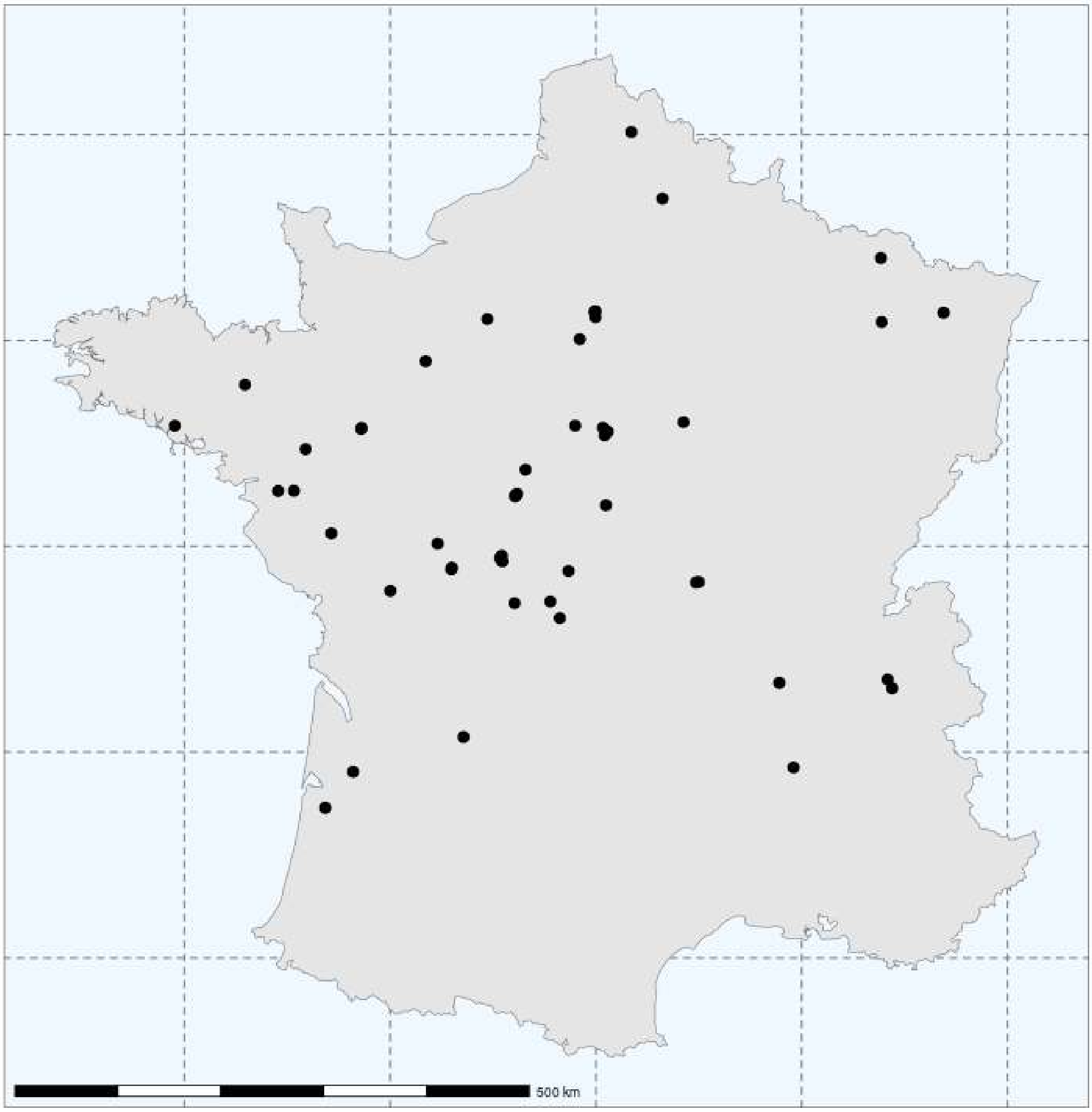

We first proceed with the spatial analysis of this point pattern, then we consider the time distribution of these cases, and finally we analyse the space-time distribution of cases.

#### A.2 Spatial analysis of the bird occurrences

##### A.2.1 K function and the complete spatial randomness

As noted in the paper, we first calculate the L function to identify scales of clustering in the spatial distribution of dead birds (Baddeley *et al.*, 2016, p. 203 and 207). We do not account for the species of the birds in this analysis:

```
## Distances for which the K function is desired (from 1 to 250 km)
vs <- seq(1000, 250000, length=100)

## The boundary of the study area
oo <- sfFrance[[1]][[1]]

## Coordinates of the points
xy <- as.matrix(usutup[,c("x","y")])

## Calculation of the K function (divide all coordinates and distance
## by 1000: representation in km is clearer than representation in
## meters)
kh <- splancs::khat(xy/1000, oo/1000, vs/1000)

## Calculation function L
kh <- sqrt(kh/pi)-vs/1000
```

We then simulate the complete spatial randomness (Baddeley *et al.*, 2016, p. 132) 999 times to derive the envelope of the set of K functions expected under this process. WARNING: THIS CALCULATION TAKES A VERY LONG TIME !!!! Note that we have included the results of this calculation as a dataset of the package, so that the reader does not need to launch this function to reproduce further calculations:

```
## Calculation enveloppe by simulation of complete spatial randomness
## (min and max for different values of distances)
simKcsr <- splancs::Kenv.csr(nrow(xy),oo/1000,999,vs/1000,quiet=TRUE)
```

We load the results of these simulations:

```
data(simKcsr)
```

And we plot the L-function with le limits of the envelope under complete spatial randomness:

```
## transformation envelope of K into envelope of L
simKcsr$lower <- sqrt(simKcsr$lower/pi)-vs/1000
simKcsr$upper <- sqrt(simKcsr$upper/pi)-vs/1000

## Display the function L with envelope expected under CSR
plot(vs/1000, kh, ty="l",
     ylim=range(c(kh,unlist(simKcsr))), xlab="distance t (km)", ylab="L(t)")
polygon(cbind(c(vs/1000, rev(vs/1000)),
              c(simKcsr$lower,rev(simKcsr$upper))), col="grey", border = NA)
abline(h=0)
```

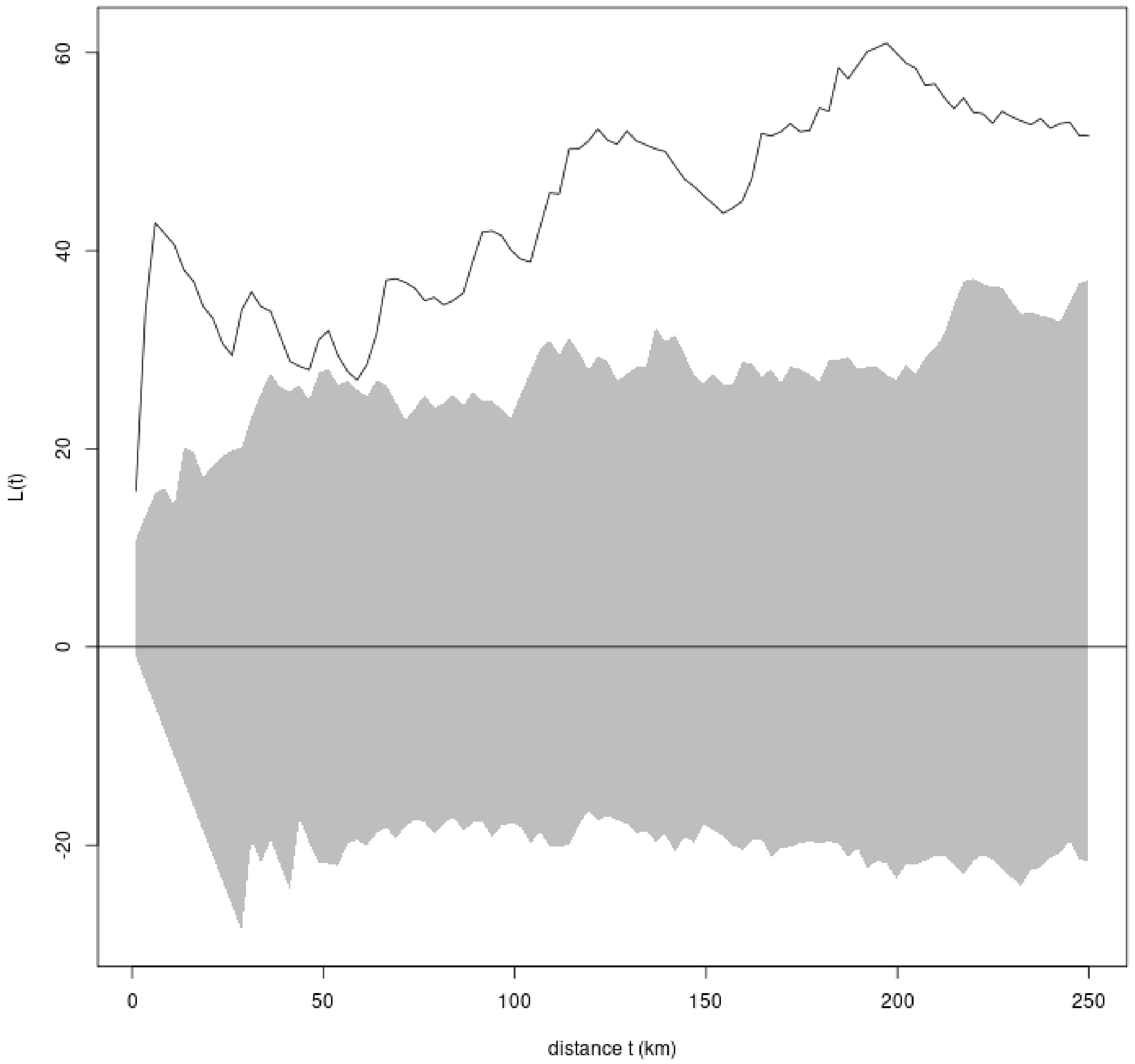

Which illustrates the clustering of the point pattern at all scales, and in particular at 5 km, 125 km and 200 km.

##### A.2.2 Fit of a Thomas process to the spatial distribution of occurrence

Given the previous result, we decided to fit a Thomas process to the location data using the minimum contrast method based on the function K (Baddeley *et al.*, 2016, p. 463). We use the functions of the package `spatstat` for this fit:

```
## Define a point pattern (similarly, divide by 1000: it is clearer to
## work in kilometers).
ppo <- spatstat.geom::ppp(xy[,1]/1000, xy[,2]/1000,
                          window =
                              spatstat.geom::owin(poly=oo[nrow(oo):1,]/1000))

## Fit the Thomas process (no trend)
ppro <- spatstat.model::kppm(ppo, ~1, "Thomas")
```

We then simulate this process 999 times to derive an envelope for the set of K functions expected under this process. WARNING: THIS CALCULATION TAKES A VERY LONG TIME !!!! Note that we have included the results of this calculation as a dataset of the package, so that the reader does not need to launch this function to reproduce further calculations:

```
simKraw <- sapply(1:999, function(r) {
    cat(r,"\r")
    s <- simulate(ppro,1)[[1]]
    khs <- splancs::khat(cbind(s$x,s$y), oo/1000, vs/1000)
    khs <- sqrt(khs/pi)-vs/1000
    return(khs)
})
```

We load the results of these simulations:

```
data(simKraw)
```

And we show the fit based on the L-function:

```
## And show the fit with:
rk <- t(apply(simKraw,1,function(x) range(x)))
vsb <- vs/1000

## prepare the plot
plot(vsb,vsb, ty="n", ylim=range(c(c(rk),kh)),
      ylab="Function L", xlab="Distance")

## show the envelope
polygon(c(vsb, rev(vsb)), c(rk[,1], rev(rk[,2])),
        col="grey", border=NA)
## and the function L
lines(vsb, kh)
```

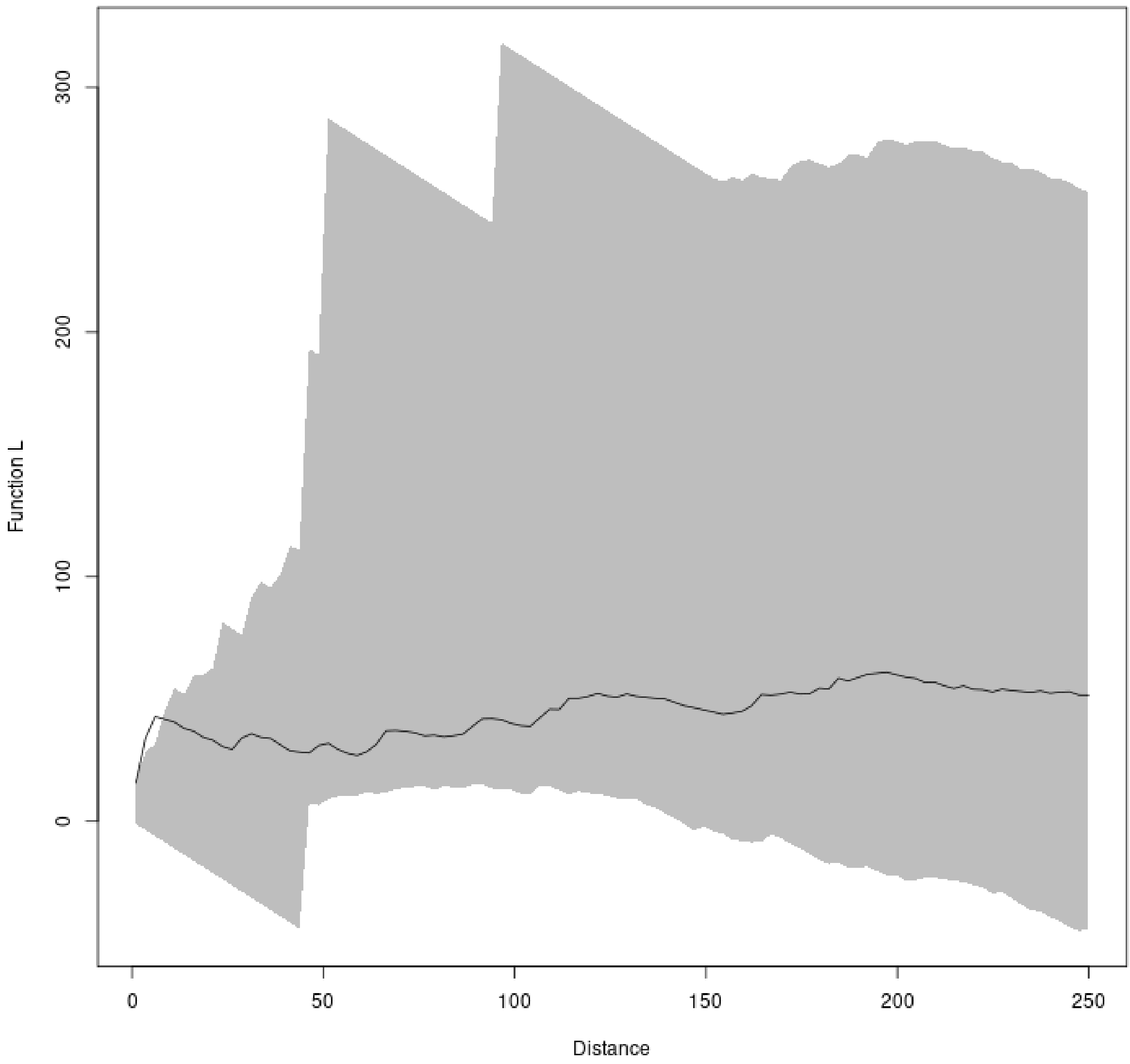

Note that at very small scales, the L-function shows a clustering not accounted for by the Thomas process. This corresponds to the small peak already identified at 5 km. This peaks corresponds to a tendency of observers to look for other dead birds in the same commune when a dead bird has been identified there by the observer.

*Remark*: One possible reason for this tendency to report several birds at the same place could be that the observers tend to search for other dead birds belonging to different species when they have reported a first dead bird nearby. If this is the case, then one would expect that animals reported at small distance from each other belong to different species more frequently than expected by chance. We carried out a randomization test to test this hypothesis (Manly, 1991). More precisely, we tested whether the pairs of birds reported at a distance lower than 5 km were more often of the same species than expected by chance. There are actually 10 pairs of birds located at less than 5 km from each others, among which 4 are from the same species:

```
## Removal of all points located at less than 5 km from another point
## Calculation of distance between points
dp <- as.matrix(dist(xy))
## identification, for eachpoint, of the index of the first point
## for which the distance is lower than 5 km (when the distance between
```

```r
## i,j is lower than 5 km, if i<j, then the first point on the i-th row
## will be the point i, and the first point on the j-th row will also be
## the point i).
firstL5 <- sapply(1:nrow(dp),function(x) min(which(dp[x,]<5000)))
## Remove duplicated points
dup <- duplicated(firstL5)

## Number of pairs:
sum(dup)

## [1] 10

## Number of pairs of the same species:
w <- which(dup)
sum(usutup$species[w]==usutup$species[w-1])

## [1] 4

## Which corresponds to two pairs blackbirds and two pairs of Great
## grey owl
usutup$species[w][usutup$species[w]==usutup$species[w-1]]

## [1] "Strix nebulosa" "Turdus merula"  "Turdus merula"
## [4] "Strix nebulosa"
```

We carried out this test by randomly permuting the species associated to the birds in our dataset and counting the number of times pairs of the same species were located at less than 5 km. We performed 999 simulations of this process. The P-value of this test is calculated below:

```r
set.seed(777)

ra <- sapply(1:999, function(r) {
    ## Random permutation
    w <- which(c(FALSE,sample(dup[-1])))
    ## Number of pairs of the same species
    sum(usutup$species[w]==usutup$species[w-1])
})

## The P-value of this test is
mean(ra<=4)

## [1] 0.4454454
```

We therefore fail to identify any significant tendency to report less birds of the same species when they are collected at the same place. Thus, the observed clustering at small distances is not clearly caused by a tendency to report birds of different species. We decided to continue to ignore the species in further analyses.

To improve the fit of the Thomas process, we decided to thin the point pattern prior to the fit, keeping only the first found bird when two birds were collected at a distance lower than 5 km:

```r
xy_thin <- xy[!dup,]
```

We then calculated the K and L functions on this thinned point pattern:

```
kh_thin <- splancs::khat(xy_thin/1000, oo/1000, vs/1000)

## Function L
kh_thin <- sqrt(kh_thin/pi)-vs/1000
```

And we fitted the Thomas process again to this thinned dataset:

```
## Define a point pattern for the package spatstat (divide by 1000 to work on km)
ppo_thin <- spatstat.geom::ppp(xy_thin[,1]/1000, xy_thin[,2]/1000,
                               window =
                               spatstat.geom::owin(poly=oo[nrow(oo):1,]/1000))

## Fit the Thomas process
ppro_thin <- spatstat.model::kppm(ppo_thin, ~1, "Thomas")
```

Again, we can simulate this Thomas process 999 times, to derive an envelope for the set of K functions expected under this process. WARNING: THIS CALCULATION TAKES A VERY LONG TIME !!!! Note that we have included the results of this calculation as a dataset of the package, so that the reader does not need to launch this function to reproduce further calculations:

```
simKthin <- sapply(1:999, function(r) {
    cat(r,"\r")
    s <- simulate(ppro_thin,1)[[1]]
    khs <- splancs::khat(cbind(s$x,s$y), oo/1000, vs/1000)
    khs <- sqrt(khs/pi)-vs/1000
    return(khs)
})
```

We load the results of these simulations:

```
data(simKthin)
```

And we show the L-function together with this calculated envelope:

```
## And show the fit with:
rk <- t(apply(simKthin,1,function(x) range(x)))
vsb <- vs/1000

## prepare the plot
plot(vsb,vsb, ty="n", ylim=range(c(c(rk),kh_thin)),
      ylab="Function L", xlab="Distance")

## show the envelope
polygon(c(vsb, rev(vsb)), c(rk[,1], rev(rk[,2])),
        col="grey", border=NA)
## and the function L
lines(vsb, kh_thin)
```

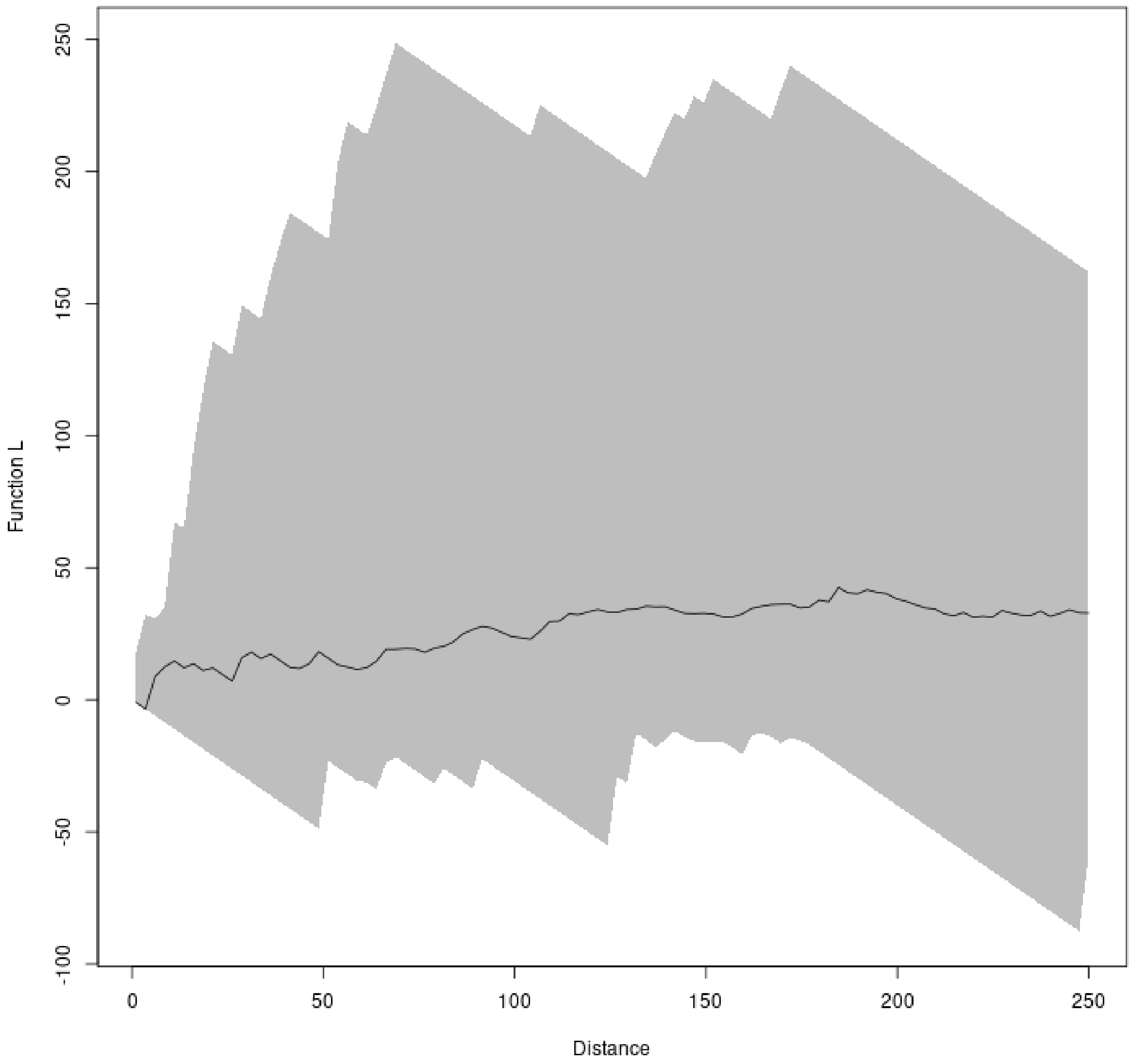

The fit is now correct. The estimated parameters of the models are therefore:

```
## The log-density of clusters
co <- coef(summary(ppro_thin))

## get the estimate and confidence limits, calculate the exponential
## to estimate the density, and multiply by 10000 to have the mean
## number of cases per 10000 squared km (mean number of cases in
## a square of 100 x 100 km)
exp(co[c(1,3,4)])*10000

##              Estimate   CI95.lo  CI95.hi
## (Intercept) 0.7408151 0.3895074 1.408977

## All the other parameters of the process
ppro_thin

## Stationary cluster point process model
## Fitted to point pattern dataset 'ppo_thin'
```

```
## Fitted by minimum contrast
##  Summary statistic: K-function
##
## Uniform intensity: 7.408151e-05
##
## Cluster model: Thomas process
## Fitted cluster parameters:
##        kappa        scale
## 1.699252e-05 7.690762e+01
## Mean cluster size:  4.359655 points
##
## Cluster strength: phi =  0.7918
## Sibling probability: psib =  0.4419
```

Thus, there is in average 0.74 cases in a square of 100×100 km (the parameter `intensity` is the density per squared km). This density of cases is the product of a density of 0.17 clusters in a square of 100 × 100 km (parameter `kappa`) and a mean cluster size of 4.36 cases per cluster. The standard deviation of the distribution of distances between the cases and the centroid of the cluster is of 77 km (parameter `scale`).

Note that this estimated standard deviation of 77 km is consistent with the K function, which identified clusters at a scale of 130 km. Indeed, this standard deviation measures the typical distance between the cases and the centroid of the cluster, whereas the K function is based on the distance between two cases. We can check by simulation that the mean distance between points sampled for a bivariate Gaussian distribution with a standard deviation of 77 km is approximately of 130 km:

```
## Simulation of 1000 points drawn from a bivariate Gaussian
## distribution with standard deviation equal to 77 km, and
## calculation of the mean distance between pairs of points.
mean(dist(cbind(rnorm(1000,0,77),rnorm(1000,0,77))))

## [1] 135.0801
```

##### A.2.3 Effect of environmental variables on the probability of a cluster

We now load the raster maps of potential wetlands and log-human population density in France (see the help page of the dataset `usutuEnvir` for more details on how these maps were built). We then show these maps, together with a kernel smoothing of the point pattern (smoothing parameter $h$=125 km, Wand and Jones, 1995), to visualize the relationship between the density of cases and the environmental variables:

```
library(sp)
data(usutuEnvir)

par(mfrow = c(1,2),mar=c(0,0,2,0))
image(usutuEnvir[1], col=viridis::magma(10))
plot(sfFrance, add=TRUE)
kd <- MASS::kde2d(xy[,1], xy[,2], 125000, n=400, lims=c(range(oo[,1]),range(oo[,2])))
contour(kd$x, kd$y,kd$z, add=TRUE, lwd=2, col="lightgrey", nlevels=4,
        levels=seq(0,1.4e-11, length=10)[c(1,4,7,10)],drawlabels = FALSE)
title("Potential Wetlands")
image(usutuEnvir[2], col=viridis::magma(10))
plot(sfFrance, add=TRUE)
contour(kd$x, kd$y,kd$z, add=TRUE, lwd=2, col="lightgrey", nlevels=4,
```

```
        levels=seq(0,1.4e-11, length=10)[c(1,4,7,10)],drawlabels = FALSE)
title("Log-Human population density")
```

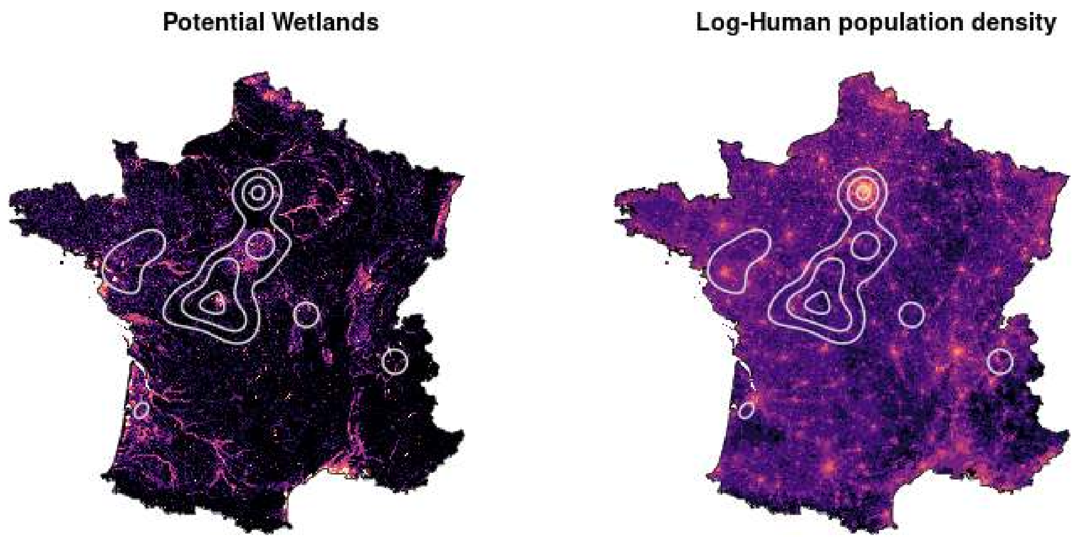

There seems to be a relationships between cases and both variables: some clusters are found in high human density areas (Paris and Nantes), whereas others are located close to wetlands (Nantes again, as well as the Brenne region – which corresponds to the main cluster at the south-west of Paris).

We used a randomization approach to test whether the mean value of the environmental variables were greater in the places where the dead birds positive to USUV were found than expected by chance (Manly, 1991). More precisely, we simulated the Thomas process fitted in the previous section 999 times over France, and we calculated for each simulation the mean of the density of humid areas and the mean of the log of human population density. We then compared these observed values with the simulated distribution. We first calculated the vector of means of these variables:

```
sppo <- sp::SpatialPoints(xy_thin, proj4string=sp::CRS(sp::proj4string(usutuEnvir)))
(obs_means <- colMeans(sp::over(sppo, usutuEnvir)))

## Wetlands  PopDens
## 4.813901 5.017741
```

The probability of wetlands is an index varying from 0 (no wetlands) to 20 (wetland certain). The population density here correspond to the mean of the log-transformed population density (transformation log(x+1), where x is expressed in number of inhabitant per squared kilometers).

We then simulate the Thomas process 999 times, and calculate the mean values of the two variables for each simulation. WARNING: THIS CALCULATION TAKES A VERY LONG TIME !!!! Note that we have included the results of this calculation as a dataset of the package, so that the reader does not need to launch this function to reproduce further calculations:

```
simEnv <- t(sapply(1:999, function(r) {
    cat(r,"\r")
    s <- simulate(ppro_thin,1)[[1]]
    sppob <- sp::SpatialPoints(cbind(s$x,s$y)*1000,
                               proj4string=sp::CRS(sp::proj4string(usutuEnvir)))
    return(colMeans(sp::over(sppob, usutuEnvir), na.rm=TRUE))
}))
```

We load the result of these simulations:

```
data(simEnv)
```

We can compare the observed means for the two variables with the simulated mean and standard error for these variables, derived from the simulations:

```
## Mean and standard error for the index of probability of wetlands
c(mean(simEnv[,1]), sd(simEnv[,1]))

## [1] 2.6915701 0.6507697

## Recall the observed mean:
obs_means[1]

## Wetlands
## 4.813901

## Mean and standard error for the log-human population density
c(mean(simEnv[,2]), sd(simEnv[,2]))

## [1] 3.5023693 0.2507175

## Recall the observed mean:
obs_means[2]

##  PopDens
## 5.017741
```

Clearly, the areas where many dead birds positive to USUV are found are characterized by a high probability of wetlands and a high density of human population.

We can display the results of this bivariate test graphically, using the function `biv.test` from the package adehabitatHS:

```
adehabitatHS::biv.test(as.data.frame(simEnv), obs_means)
```

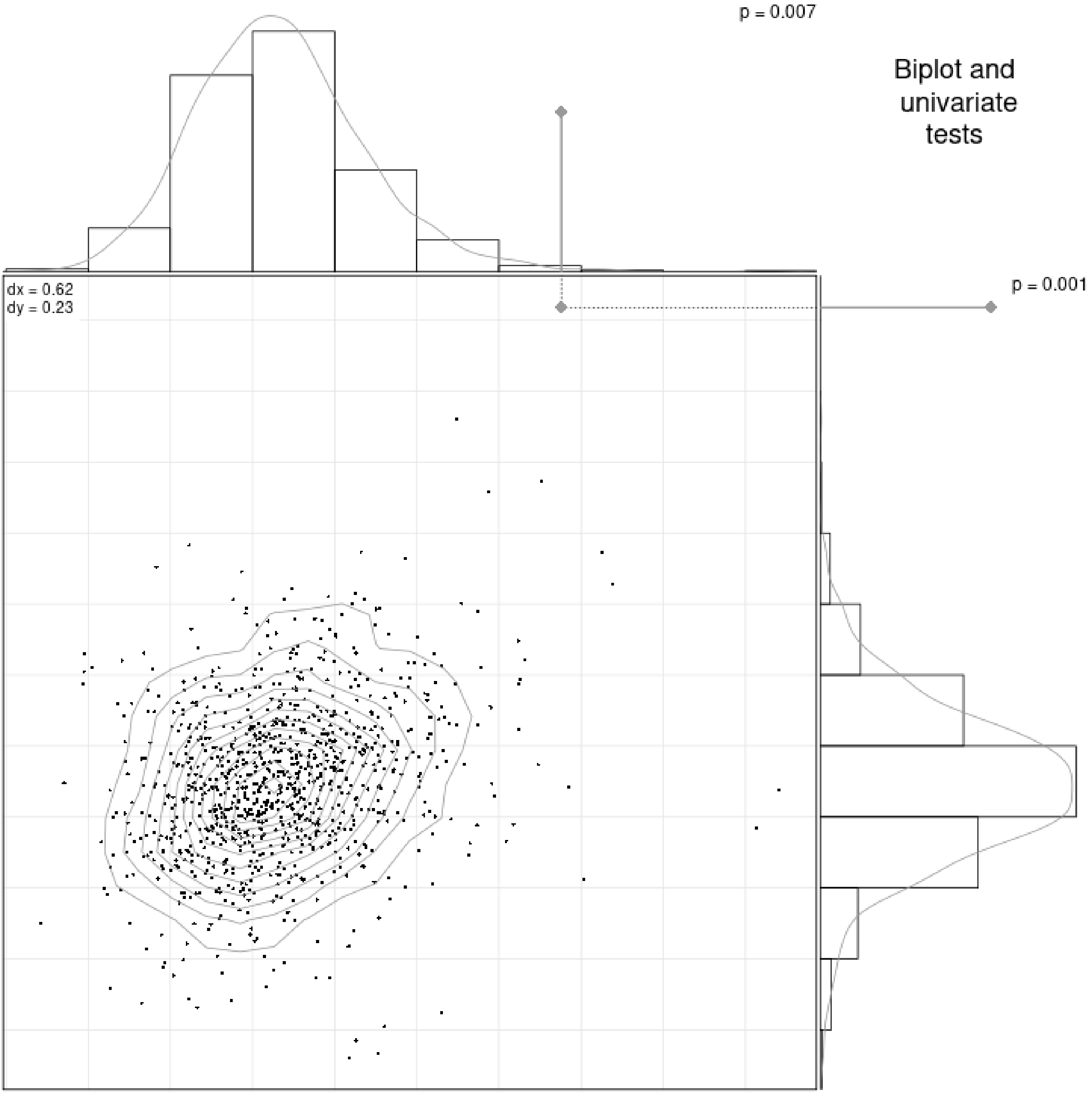

The abscissa corresponds to the probability of wetlands, and the ordinate corresponds to the log-human population density. Each small black point corresponds to the vector of means of environmental variables for one simulation of the Thomas process, whereas the grey point corresponds to the observed value for this vector of means. The margins of this graph show the statistical distribution of these means expected by chance, as well as the observed value for each variable. These margins also show the P-values of each univariate test (which are presented in the main text). This test confirms that USUV cases are more frequent in places where both human population and wetlands are higher.

#### A.3 Temporal and Spatio-temporal distribution of bird occurrences

We now present how the mean number of cases per day change throughout the study period. We smoothed this curve using moving average approach (Diggle, 1990, p. 22): we present, for each day $d$ of the study period, the mean number of cases per day calculated over a week centred on $d$:

```
library(lubridate)
seqPeriod <- seq(ymd('2018-07-15'),ymd('2018-08-31'), by = '1 day')
movAvg <- sapply(1:length(seqPeriod),  function(i) {
    x <- seqPeriod[i]
```

```
    ## days of the week centred on the day i
    d <- seq(x-3, x+3, by="1 day")
    ## number of days of the week in the study period
    ndays <- mean(d>=seqPeriod[1]&d<=seqPeriod[length(seqPeriod)])
    ## number of cases reported in this period
    ncases <- sum(usutup$date>=(seqPeriod[i]-3)&usutup$date<=(seqPeriod[i]+3))
    ## number of cases per day:
    return(ncases/ndays)
})

da <- data.frame(date=seqPeriod,
                 NumberCases=movAvg)

ggplot(da, aes(x=date,y=NumberCases))+geom_line()+xlab("Date")+
    ylab("Mean number of cases reported per day")
```

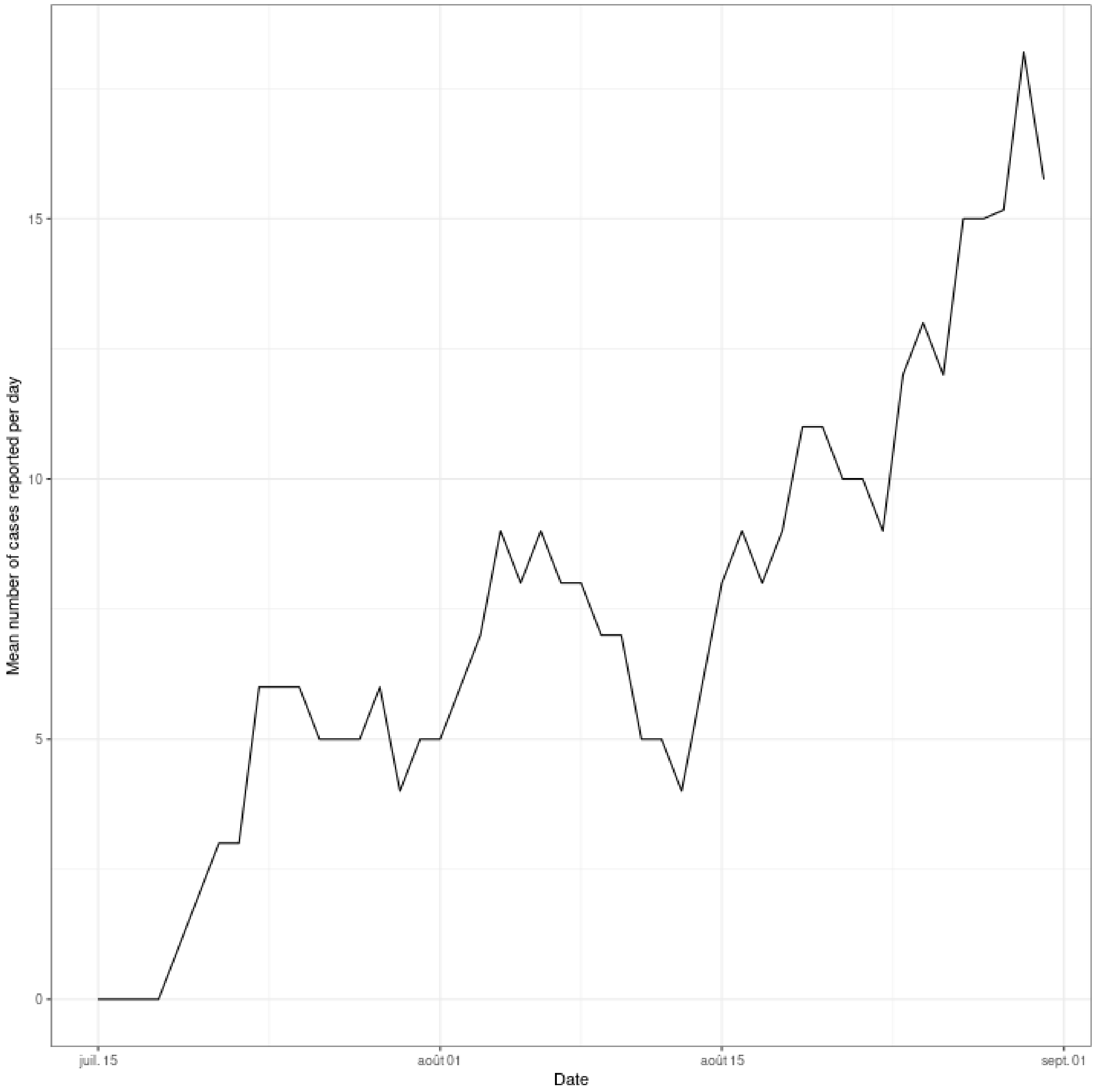

A graphical analysis of the spatio-temporal patterns in USUV occurrence starts with a map of the occurrence where the color indicates the time at which the bird was found:

```r
theme_set(theme_bw())
labd <- pretty(usutup$date)
ggplot()+
    geom_sf(data=sfFrance)+
    geom_point(aes(x=x,y=y,col=as.numeric(date),group=as.factor(date)),data=usutup,size=3)+
    theme(panel.grid.major = element_line(color = gray(0.5),
                                          linetype = "dashed",
                                          linewidth = 0.5),
          panel.background = element_rect(fill = "aliceblue"))+
    theme(axis.line=element_blank(),
          axis.text.x=element_blank(),
          axis.text.y=element_blank(),
          axis.ticks=element_blank(),
          axis.title.x=element_blank(),
          axis.title.y=element_blank(),
          legend.position="none")+
    viridis::scale_color_viridis(breaks = as.numeric(labd),
                                 labels = labd)+
    annotation_scale(location = "bl", width_hint = 0.5, plot_unit="m")
```

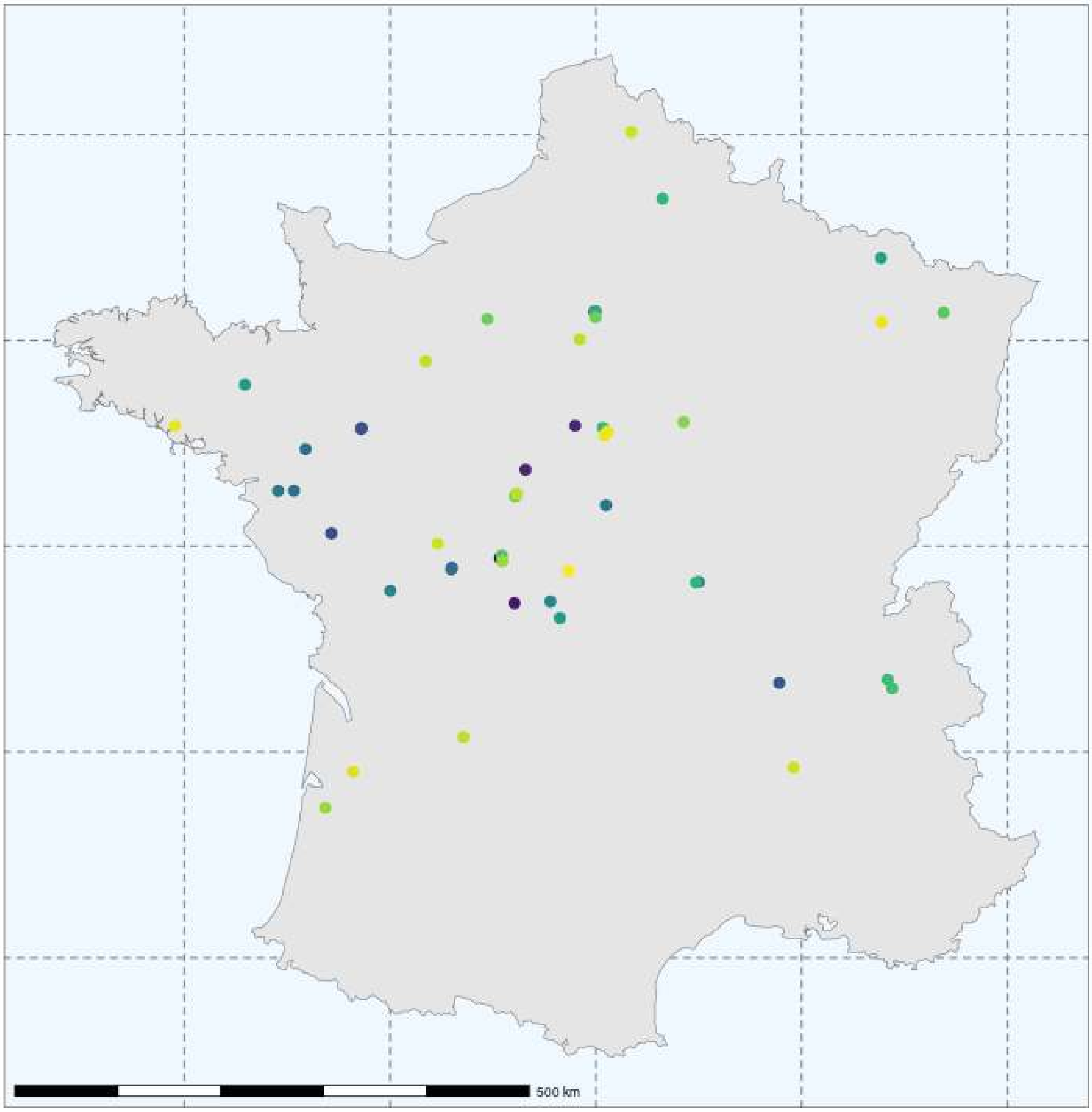

Blue points correspond to birds found at mid-July, whereas yellow points correspond to birds reported by the end of August. It seems that the main clusters from center and western France appeared approximately at the same time at about mid-July.

We calculated the Space-Time K-function based on this point pattern (Gabriel *et al.*, 2013):

```
## get define space-time points
po <- stpp::as.3dpoints(usutup$x/1000,usutup$y/1000,
                        as.numeric(usutup$date-min(usutup$date)))

## The set of distances for which the function K is calculated
## (between 1 to 220 km)
vs <- seq(1000, 220000, length=100)
## The weeks
vt <- 1:10

## the dates
dates <- as.numeric(usutup$date-min(usutup$date))
```

```r
## Space-time function
ch <- stpp::STIKhat(po, s.region = oo/1000,
                    t.region = c(min(dates-1), max(dates+1)),
                    times=vt, dist=vs/1000, infectious=TRUE)
```

We then wanted to know whether birds found at small distance from each other were also found at close dates. We carried out a set of simulations to test whether this space-time K function was greater than expected by chance, by randomizing the collection dates associated with the bird locations. WARNING: THIS CALCULATION TAKES A VERY LONG TIME !!!! Note that we have included the results of this calculation as a dataset of the package, so that the reader does not need to launch this function to reproduce further calculations:

```r
## Randomization test: WARNING! THIS PART IS VERY SLOW
si <- list()

## 999 randomization (slow calculation !!)
for (i in 1:999) {
    cat(i,"\r")
    pos <- po
    pos <- stpp::as.3dpoints(usutup$x/1000, usutup$y/1000,sample(dates))
    chs <- stpp::STIKhat(pos, s.region = oo/1000,
                         t.region = c(min(dates-1), max(dates+1)),
                         times=vt, dist=vs/1000, infectious=TRUE)
    si[[i]] <- chs$Khat
}

## Calculation of the proportion of simulations greater than observation
KstFun <- ch$Khat ## Initialization
for (i in 1:nrow(KstFun)) {
    cat(i,"\r")
    for (j in 1:ncol(KstFun)) {
        ot <- sapply(1:length(si), function(k) si[[k]][i,j])
        KstFun[i,j] <- mean(ot>ch$Khat[i,j])
    }
}
```

We load the dataset containing the resulting matrix:

```r
data(KstFun)
```

We then show, for each distance and duration between occurrence, the probability that the observed space-time K function is greater than the set of functions simulated under the hypothesis of absence of time structure:

```r
## Show the results:
dok <- data.frame(duration=rep(vt,length(vs)),
                  distance=rep(vs/1000,each=length(vt)),
                  Prob. =as.vector(t(KstFun)))

ggplot2::ggplot(dok, ggplot2::aes(x=duration, y=distance))+
    ggplot2::geom_tile(ggplot2::aes(fill=Prob.))+
    viridis::scale_fill_viridis(option="magma", limits=c(0,1))+
    ggplot2::geom_contour(ggplot2::aes(z=Prob.), breaks = 0.05, col="white",
                          alpha=0.5, linewidth=1)+ggplot2::theme_bw()+
    xlab("Duration (days)")+ylab("Distance (km)")
```

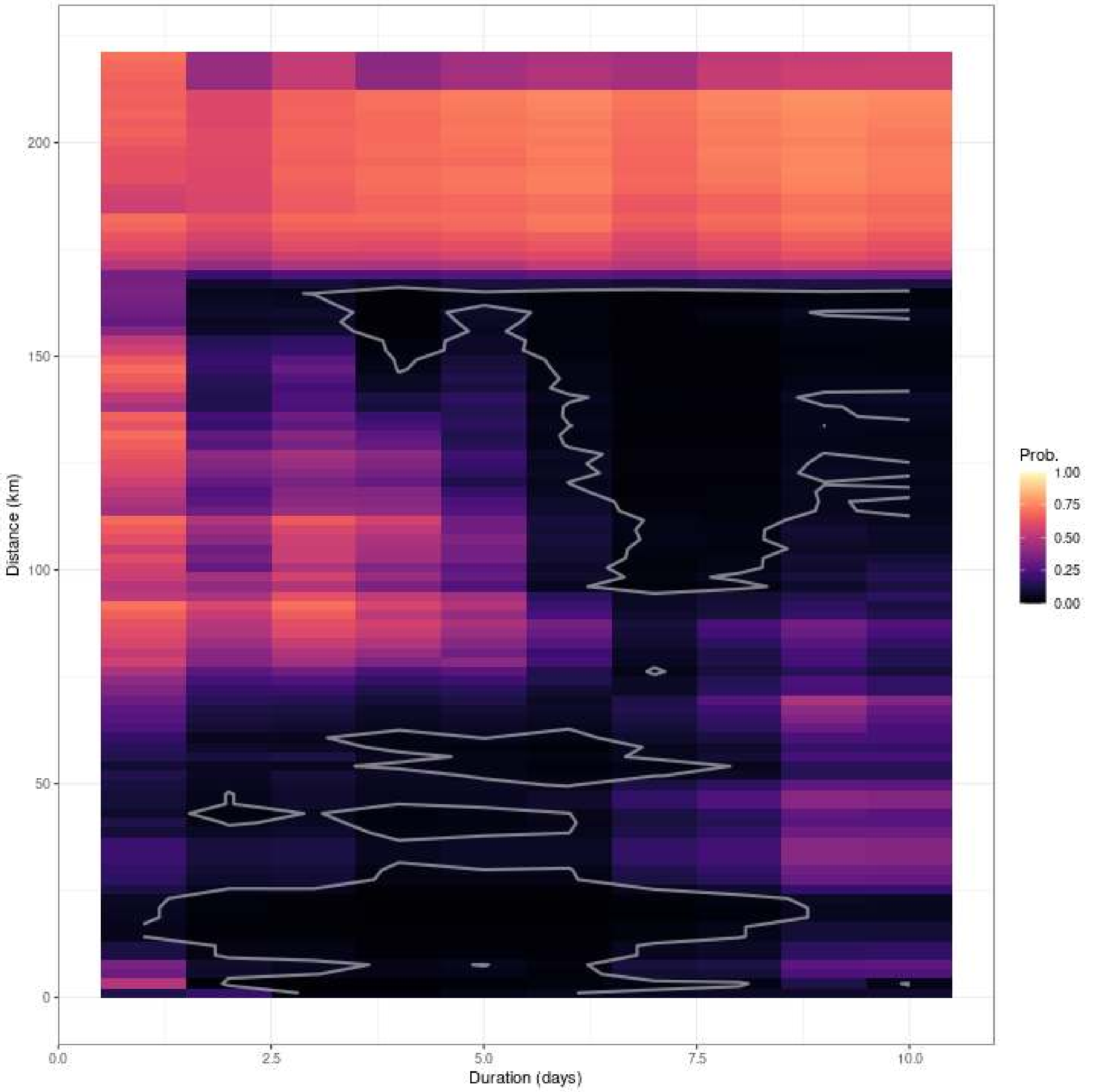

The contour limit identify the set of distances/duration pairs for which the proportion of simulated K functions greater than the observed K function is lower than 5%.

There is a strong clustering for small distances (as soon as 2 or 3 days after a case) of about 10 to 20 km, expanding to 50-60 km five days after a case. But we also observe a clustering on larger distances (150 km) starting as soon as 2 or 3 days after a case, which increases to reach a large range of distances after a week (90 to 150 km).

#### A.4 Synthesis figure

We finally draw the synthesis figure that is displayed in the main paper (Fig. 1), and which summarizes the main results of this analysis:

```
##png(filename="SynthesisFigure.png", width=1000, height=1000, pointsize=20)
library(gridBase)
library(grid)

par(mfrow=c(2, 2))
```

```
plot(vs/1000, kh, ty="l",
     ylim=range(c(kh,unlist(simKcsr))), xlab="Distance t (km)", ylab="L(t)",
     main="(A)")
polygon(cbind(c(vs/1000, rev(vs/1000)),
              c(simKcsr$lower,rev(simKcsr$upper))), col="grey", border = NA)
abline(h=0)

opar <- par(mar=c(1,1,4,1))
kd <- MASS::kde2d(xy[,1], xy[,2], 125000, n=400, lims=c(range(oo[,1]),range(oo[,2])))
plot(sfFrance, main="(B)", col=grey(0.8), border=NA)
contour(kd$x, kd$y,kd$z, add=TRUE, lwd=2, col=grey(0.6), nlevels=4,
        levels=seq(0,1.4e-11, length=10)[c(1,4,7,10)],drawlabels = FALSE)
usud <- unclass(usutup$date-min(usutup$date))
points(usutup[,c("x","y")], pch=3, col="black", cex=1.5)

da <- data.frame(date=seqPeriod,
                 NumberCases=movAvg)
par(opar)
plot(da$date, da$NumberCases, ty="l",xlab = "Date",
     ylab="Mean number of cases reported per day", main="(C)")
plot.new()
vps <- baseViewports()
pushViewport(vps$figure)
vp1 <-plotViewport(c(1.8,1,3,1))
p <- ggplot2::ggplot(dok, ggplot2::aes(x=duration, y=distance))+
    ggplot2::geom_tile(ggplot2::aes(fill=Prob.))+
    viridis::scale_fill_viridis(option="magma", limits=c(0,1))+
    ggplot2::geom_contour(ggplot2::aes(z=Prob.), breaks = 0.05, col="white",
                          alpha=0.5, linewidth=0.8)+ggplot2::theme_bw()+
    theme(axis.text=element_text(size=12),
          axis.title=element_text(size=13))+
    xlab("Duration (days)")+ylab("Distance (km)")+title("(D)")
print(p,vp = vp1)
##dev.off()
```

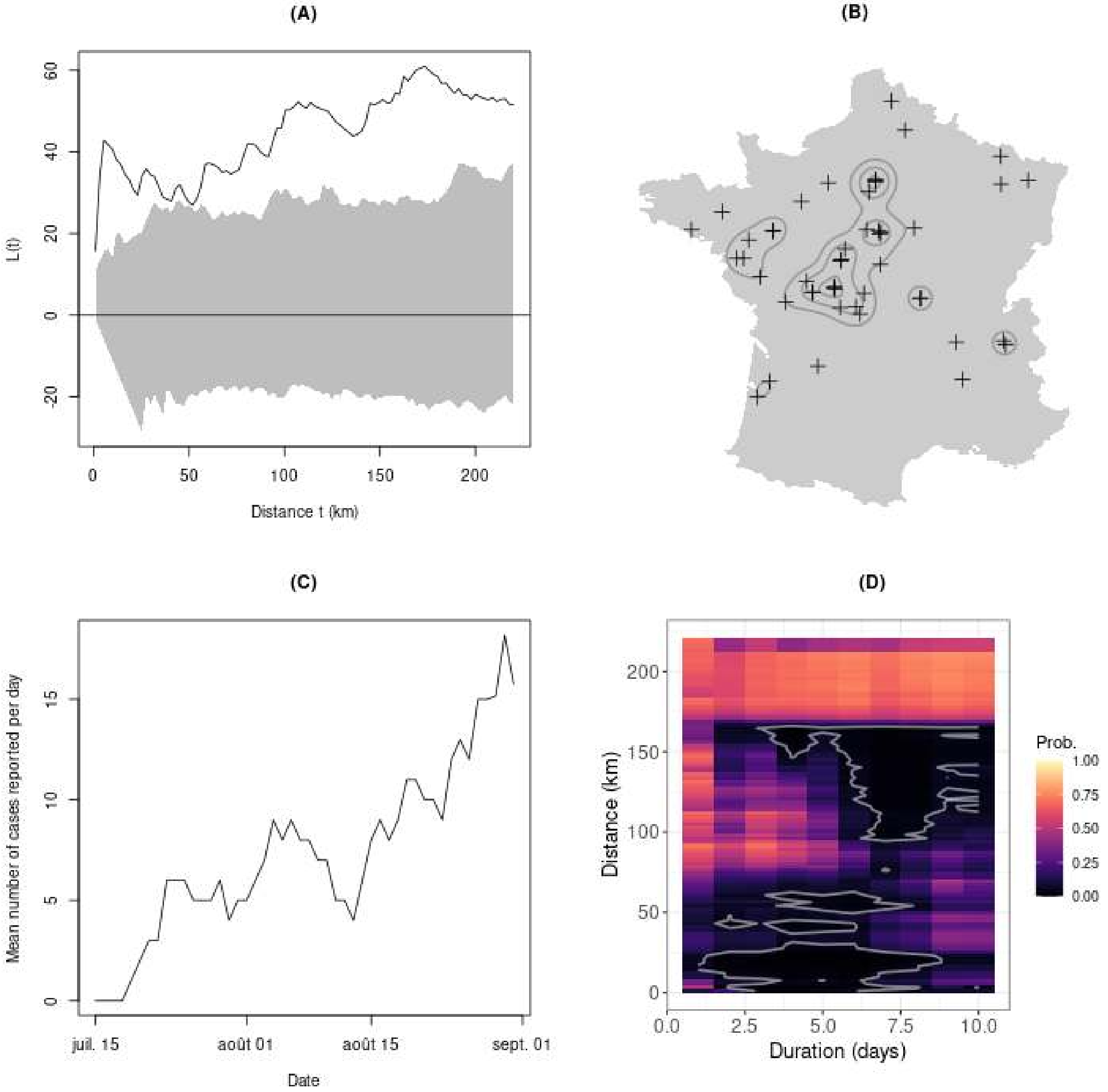

(A)
L(t)
60
40
20
0
-20
0
50
100
150
200
Distance t (km)
(B)
(C)
Mean number of cases reported per day
15
10
5
0
juil. 15
août 01
août 15
sept. 01
Date
(D)
Distance (km)
200
150
100
50
0
0.0
2.5
5.0
7.5
10.0
Duration (days)
Prob.
1.00
0.75
0.50
0.25
0.00

### Appendix B  Assessment of the population trends as a function of the estimated level of infection

In this appendix, we provide the R code used to correlate the level of infection by the Usutu virus in a region with the changes in blackbird population size in this region.

#### B.1  Definition of the regions with the three levels of infection

We will need the object `kd` created in appendix A, which stores the density of cases over continental France, estimated by the kernel smoothing method. We first convert this object into an object of class `spatRaster` from the package `terra`:

```
## Load the package terra
library(terra)

## Convert to spatRaster
m <- kd
resx <- diff(kd$x)[1]
resy <- diff(kd$y)[1]
xmn <- min(m$x) - 0.5 * resx
xmx <- max(m$x) + 0.5 * resx
ymn <- min(m$y) - 0.5 * resy
ymx <- max(m$y) + 0.5 * resy
z <- m$z[,ncol(m$z):1]
r1 <- rast(ext(xmn, xmx, ymn, ymx), resolution=c(resx, resy))
r1[] <- as.numeric((z))

## Conversion of sfFrance to spatVector
sfv <- vect(sfFrance)

## Set the map to NA outside the limits of continental France
rr <- rasterize(sfv, r1)
r1 <- rr*r1

image(r1)
```

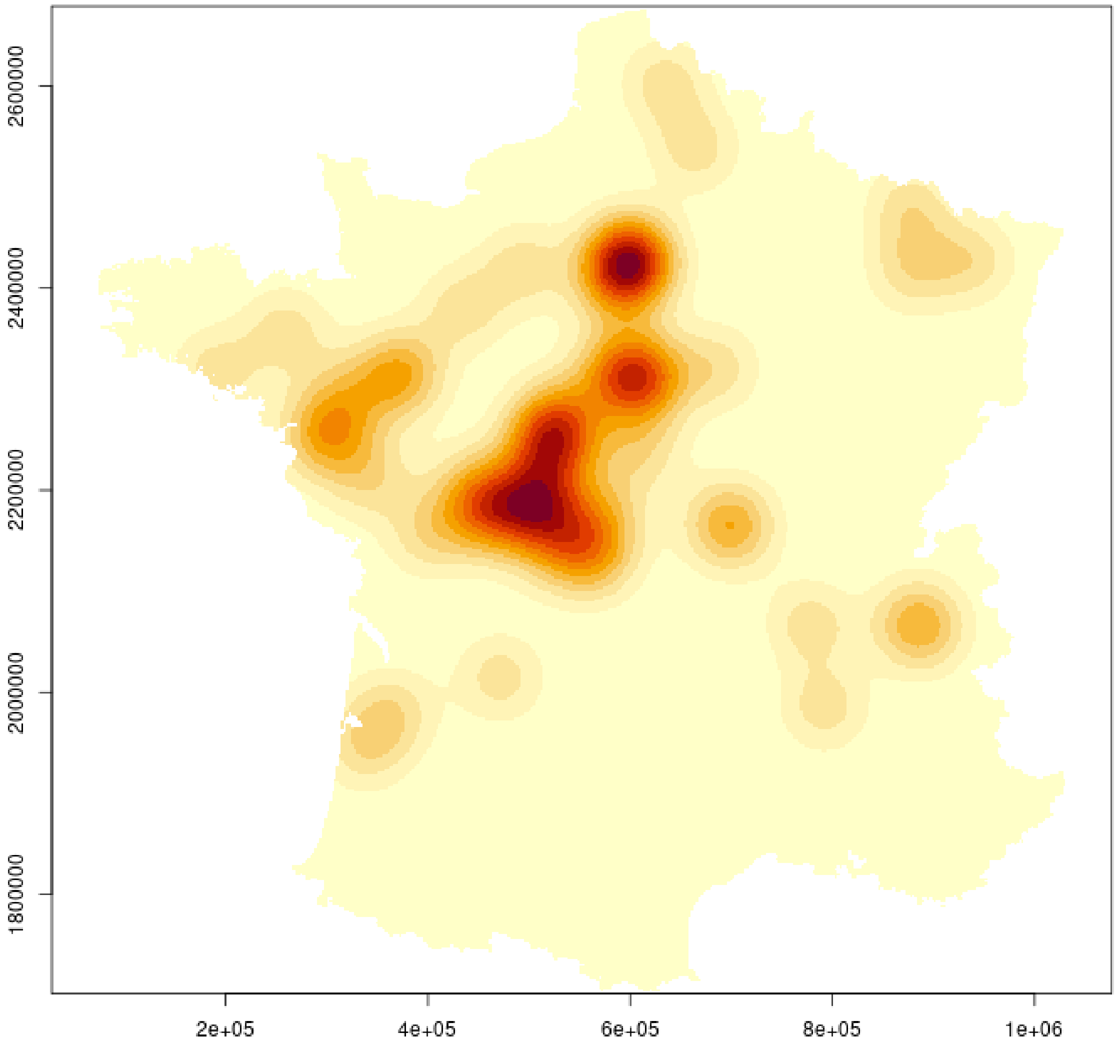

Now, we load the location of the sampling routes and points in the ACT survey, stored in the dataset `resop`:

```
data(resop)
head(resop)

##   id_point cd_route no_point cd_route_point insee_dep
## 1       38     1015        3         1015_3        22
## 2       39     1016        3         1016_3        22
## 3       40     1017        3         1017_3        22
## 4       41     1018        3         1018_3        56
## 5       42     1019        3         1019_3        56
## 6       43     1020        3         1020_3        56
##          x       y
## 1 255180.0 2415340
## 2 255380.0 2403005
## 3 253523.8 2378385
## 4 247617.0 2359967
## 5 252360.0 2341680
```

```
## 6 250360.0 2322560

plot(resop[,c("x","y")], asp=1)
```

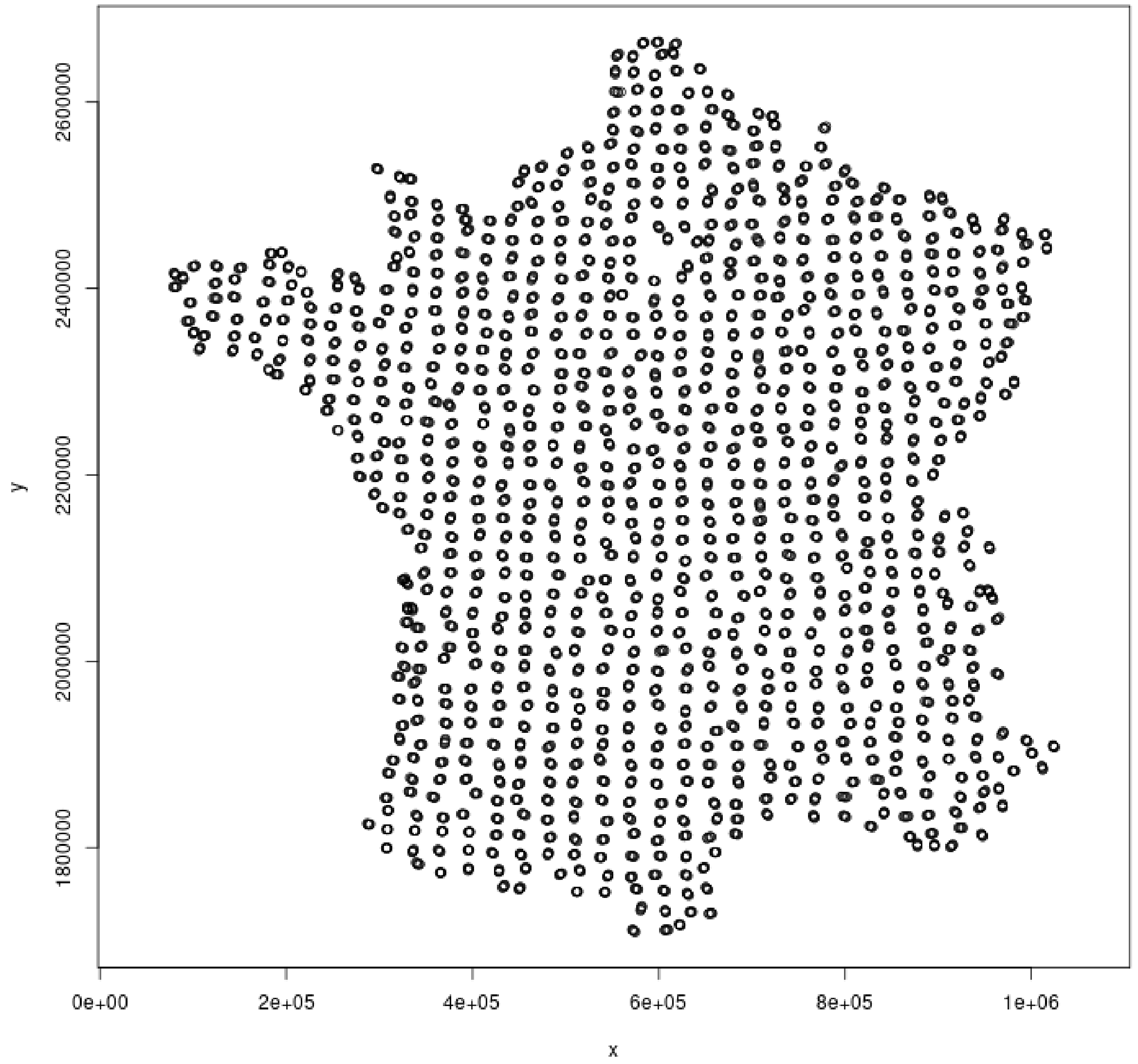

We then join these locations to the map of the USUV density, and store the result in the object `resop`:

```
resop$usutu<-terra::extract(r1, as.matrix(resop[,c("x", "y")]))[,1]
```

We then average the USUV density per route (calculation of the mean density over all sampling points of a route, as well as calculation of the centoid of the route):

```
val_agg <- resop  |>  dplyr::group_by(cd_route) |>
dplyr::summarise(usutu_agg=mean(usutu, na.rm=T),
                 x=mean(x), y=mean(y))
head(val_agg)

## # A tibble: 6 x 4
```

```
##   cd_route usutu_agg        x         y
##   <chr>         <dbl>    <dbl>     <dbl>
## 1 1015      5.73e-13 255204. 2415061.
## 2 1016      1.10e-12 255307. 2402961.
## 3 1017      2.59e-12 253418. 2378671.
## 4 1018      3.30e-12 247564. 2359936.
## 5 1019      3.32e-12 252242. 2341596.
## 6 1020      2.66e-12 250260. 2322750.
```

We then calculate the median and third quartile of the density distribution. These density values will serve to define three levels of infections: low (areas with density lower than the median), medium (areas with density comprised between the median and the third quartile), and high (areas with density greater than the third quartile):

```
## Median
usutu_med <- median(val_agg$usutu_agg)

## Third quartile
usutu_75 <- quantile(val_agg$usutu_agg,  probs=0.75)

## new variable in val_agg, defining the three levels of infection
val_agg$usutuF <- ifelse(val_agg$usutu_agg<=usutu_med, "low",
                  ifelse(val_agg$usutu_agg>usutu_75, "high", "medium"))
val_agg$usutuF <- factor(val_agg$usutuF, levels=c("low", "medium", "high"))

head(val_agg)

## # A tibble: 6 x 5
##   cd_route usutu_agg        x         y usutuF
##   <chr>         <dbl>    <dbl>     <dbl> <fct>
## 1 1015      5.73e-13 255204. 2415061. medium
## 2 1016      1.10e-12 255307. 2402961. medium
## 3 1017      2.59e-12 253418. 2378671. high
## 4 1018      3.30e-12 247564. 2359936. high
## 5 1019      3.32e-12 252242. 2341596. high
## 6 1020      2.66e-12 250260. 2322750. high
```

#### B.2 Formatting the results of ACT survey

Now, we load the results of the counts of singing male blackbirds for all routes between 1996 and 2022, which are stored in the dataset `merula`:

```
data(merula)
head(merula)

##   tot_MN        x       y            dateTime year
## 1      6 357582.0 2315954 1996-04-01 07:22:00 1996
## 2     12 605490.0 1753844 1996-04-01 07:00:00 1996
## 4      7 403730.0 2073529 1996-04-02 07:05:00 1996
## 5      1 849576.0 2115168 1996-04-02 08:00:00 1996
## 6      0 865403.2 1833320 1996-04-02 07:30:00 1996
## 7      5 938164.4 1974468 1996-04-02 07:15:00 1996
##          TsSR cd_routeF yearF jd passageN
## 1 -20.233333      1420   1996 92    First
## 2 -32.883333      2348   1996 92    First
```

```
## 4 -34.216667      1632  1996 93    First
## 5  44.016667      3230  1996 93    First
## 6  12.183333      3244  1996 93    First
## 7   2.116667      3537  1996 93    First
```

The meaning of the columns of this data.frame is explained on the help page of `merula`. We join this dataset and the raster map `r1` storing the density of usutu cases, and we store the results in the dataset `merula`. We also define a factor containing the three levels of infections (based on the limits calculated before):

```
## Join with the density map
ex2 <- terra::extract(r1, as.matrix(merula[,c("x","y")]))

## Store the results in the dataset:
merula$usutu_agg <- ex2[,1]

## median and third quartile limits:
limi <- c(usutu_med, usutu_75)

## Cut the density in three classes and store the result in merula:
cu <- cut(merula$usutu_agg, c(-10,limi,10))
cu <- factor(cu, labels=c("low","medium","high"))
merula$usutuF <- cu
```

#### B.3 Model fit and prediction

We now fit the Generalized Additive Mixed Model (GAMM) to predict the count outcome as a function of year, julian date, time since sunrise, sampling route, and the level of infection by the Usutu virus (see main text for an explanation). WARNING: THIS CALCULATION TAKES A VERY LONG TIME (2-3 hours on a computer equipped with Intel Core I7) !!!! The results of this fit, which occupy a large space (52 MB), are not included in the package. After a check of the model fit, we used the function `compute_distri` from our package to simulate 1000 count predictions for each year between 1996 and 2022 on an average road located in regions characterized by varying levels of Usutu virus infection – low, medium, or high. More precisely, 1000 coefficient vectors are simulated from a multivariate Gaussian distribution using GAMM results (with distribution parameters corresponding to the mean coefficient vector and the covariance matrix of coefficients extracted from the model). Each simulated coefficient vector is used to predict the response variable (total number of male blackbirds counted on an average route) for each year in each of the three regions. These 1000 simulations per region allow the assessment of prediction uncertainty. Note that we have included the results of this simulation as a dataset of the package, so that the reader does not need to execute this function to reproduce further calculations:

```
## Load the package mgcv
library(mgcv)

## Model fit with bam (large dataset)
m_per <- bam(tot_MN ~  usutuF + s(year, k=10) +  s(year, by=usutuF, k=10) +
                 s(cd_routeF, bs="re") + s(yearF, bs="re") +
                 s(jd, k=6) + s(TsSR, k=6) , family=tw(),
             data=merula, method="REML")

## Check the model fit
gratia::appraise(m_per)
## Nothing worrying there

## Summary of the model
```

```r
summary(m_per)

## Plot the smooth functions of explanatory variables:
plot(m_per)

## A data.frame for predictions, with unknown levels for the 2 random effects factors,
## to compute changes independently of these effects
df_pred<-data.frame(expand.grid(usutuF=levels(merula$usutuF), cd_routeF="1",
                                year=1996:2022, jd=mean(merula$jd),
                                TsSR=mean(merula$TsSR), yearF=as.factor(2030)))

#Load the functions to compute simulations and estimate change
simu_by_regions<-compute_distri(m_per, df_pred, time_var="year",
                                factor_name="usutuF", nreplicates=1000)
```

We load the resulting dataset:

```r
data(simu_by_regions)
str(simu_by_regions)

## List of 3
##  $ low   : num [1:1000, 1:27] 6.15 5.74 6.23 6.2 5.93 ...
##   ..- attr(*, "dimnames")=List of 2
##   .. ..$ : NULL
##   .. ..$ : chr [1:27] "1996" "1997" "1998" "1999" ...
##  $ medium: num [1:1000, 1:27] 7.14 6.7 7.09 7.5 7.47 ...
##   ..- attr(*, "dimnames")=List of 2
##   .. ..$ : NULL
##   .. ..$ : chr [1:27] "1996" "1997" "1998" "1999" ...
##  $ high  : num [1:1000, 1:27] 7.02 6.58 6.85 6.58 6.87 ...
##   ..- attr(*, "dimnames")=List of 2
##   .. ..$ : NULL
##   .. ..$ : chr [1:27] "1996" "1997" "1998" "1999" ...
```

See the help page of this dataset for more details. We transform this list into a data.frame (layout: one simulation per row, corresponding to a unique combination of simulation ID, infection level, and year):

```r
#convert to a data frame from which to average values per relevant time periods
df_4analyses<- do.call("rbind", simu_by_regions) |> as.data.frame() |>
dplyr::mutate(usutuF=rep(levels(merula$usutuF), each=1000),
       id_sim=rep(1:1000, 3))  |>
tidyr::pivot_longer(!c(usutuF, id_sim), names_to="year",
                    values_to="abundance")  |>
dplyr::mutate_at( 'year', as.numeric)

head(df_4analyses)

## # A tibble: 6 x 4
##   usutuF id_sim  year abundance
##   <chr>   <int> <dbl>     <dbl>
## 1 low         1  1996      6.15
## 2 low         1  1997      6.17
## 3 low         1  1998      6.19
## 4 low         1  1999      6.20
## 5 low         1  2000      6.22
## 6 low         1  2001      6.24
```

#### B.4 Calculation of trends

We now calculate, for each simulation and each infection level, the average abundance over the years preceding the usutu infection (ranging between 2015 and 2018) and the years following this infection (ranging between 2019-2022). This comparison will enable to assess the differences in mean abundance before and after the infection:

```
## A data frame where abundance is averaged by simulation,
## by period (before after the 2018 Usutu episode) and
## by areas of Usutu "prevalence" (usutuF)
df_4trends<- df_4analyses  |>
dplyr::filter(year>2014)  |>
dplyr::mutate(usutu_BA=factor(ifelse(year<2019, "2015-2018", "2019-2022"),
                                levels=c("2015-2018", "2019-2022")))  |>
dplyr::group_by(id_sim, usutu_BA, usutuF)  |>
dplyr::summarise(mean_abund=mean(abundance))

## `summarise()` has grouped output by 'id_sim', 'usutu_BA'. You can override using the `.groups`
argument.

## Define the order of the levels of infection
df_4trends$usutuF<-factor(df_4trends$usutuF, levels=c("low", "medium", "high"))

#Compute the trends for the 3 areas according to the previous values
df_trends<- df_4trends  |>
dplyr::group_by(id_sim, usutu_BA, usutuF) |>
tidyr::pivot_wider(names_from=usutu_BA, values_from=mean_abund) |>
as.data.frame()

head(df_trends)

##   id_sim usutuF 2015-2018 2019-2022
## 1      1   high  6.707626  5.815715
## 2      1    low  6.540335  6.614478
## 3      1 medium  6.898128  6.519382
## 4      2   high  6.232447  5.316868
## 5      2    low  6.111560  6.195160
## 6      2 medium  6.676841  6.234477
```

We can calculate the rate of variation $100 \times (X_2 - X_1)/X_1$ for each simulation and each level of infection (with $X_1, X_2$ the predicted count respectively before and after the infection). Then, we average the 1000 rates of variations for each level of infection and we calculate the 95% confidence interval:

```
## Low level of infection:
dl <- df_trends[df_trends$usutuF=="low",]
trl <- 100*(dl[,4]-dl[,3])/dl[,3]
tr_low <- c(mean(trl), quantile(trl, c(0.025,0.975)))

## Medium level of infection:
dm <- df_trends[df_trends$usutuF=="medium",]
trm <- 100*(dm[,4]-dm[,3])/dm[,3]
tr_medium <- c(mean(trm), quantile(trm, c(0.025,0.975)))

## High level of infection:
dh <- df_trends[df_trends$usutuF=="high",]
trh <- 100*(dh[,4]-dh[,3])/dh[,3]
tr_high <- c(mean(trh), quantile(trh, c(0.025,0.975)))
```

```r
#the final data.frame which can be turned into a table for the MS
res_trends<-data.frame(UsutuArea=c("low", "medium", "high"),
                       as.data.frame(rbind(tr_low, tr_medium, tr_high)))
colnames(res_trends)<-c("Usutu area", "Mean trend", "2.5%CI", "97.5%CI")
rownames(res_trends)<-NULL

## Results:
res_trends

##   Usutu area  Mean trend      2.5%CI   97.5%CI
## 1        low   0.7176586  -0.3102161  1.748856
## 2     medium  -7.3722991 -11.1112177 -3.669769
## 3       high -12.7091388 -16.0061863 -9.122092
```

These values correspond to those presented in the results section of the paper.

#### B.5 Summary plot

We finally build the summary graph, corresponding to Fig. 2 of the paper:

```r
## format the data.frame df_4trends for an easier handling
df_stat_abund<-df_4trends  |>
dplyr::group_by(usutu_BA, usutuF)  |>
dplyr::summarise(averag_abund=mean(mean_abund),
                 CI_low=quantile(mean_abund, probs=0.025),
                 CI_high=quantile(mean_abund, probs=0.975))  |>
dplyr::mutate_at(c('CI_low', 'CI_high'), as.numeric)

## `summarise()` has grouped output by 'usutu_BA'. You can override using the `.groups`
argument.

## correspondance between Usutu areas and colours
## for the following plots (scale_colour_manual)
col_low<-"palegreen4"
col_medium<-"darkorange2"
col_high<-"deeppink4"
val_colours<-c(col_low, col_medium, col_high)

## MAIN PLOT
## A plot of each simulated average values of abundance before and
## after, and the mean and 95%CI for each area
p_trends <- ggplot2::ggplot(df_4trends) +
    ggplot2::geom_line(ggplot2::aes(x=usutu_BA,  y=mean_abund,
                  group=id_sim, colour=usutuF), alpha=0.075) +
    ggplot2::geom_point(data=df_stat_abund,
                        ggplot2::aes(x=usutu_BA, y=averag_abund,
                                     colour=usutuF), size=4) +
    ggplot2::geom_errorbar(data=df_stat_abund,
                           ggplot2::aes(x=usutu_BA, ymin=CI_low, ymax=CI_high,
                                        colour=usutuF),  width=0.1, linewidth=1) +
    ggplot2::scale_colour_manual(name= "Usutu area", values=val_colours)+
    ggplot2::xlab("Period") + ggplot2::ylab("Abundance index") +
    ggplot2::facet_wrap(~usutuF) +
    ggplot2::theme( strip.text.x = ggplot2::element_blank() )
```

```r
## Histogram of Usutu densities values for the centroid of each route
## and categorisation in 3 groups
usutu_histo <- ggplot2::ggplot(val_agg) +
    ggplot2::geom_histogram(ggplot2::aes(x=usutu_agg, fill=usutuF)) +
    ggplot2::geom_vline(xintercept=c(usutu_med, usutu_75),
                        colour="black", linewidth=1,
                        linetype=c("solid", "dashed")) +
    ggplot2::ylab("Count") + ggplot2::xlab("Density of Usutu cases") +
    ggplot2::scale_fill_manual(name="Usutu area", values=val_colours)

## Spatial plot of ACT and density of Usutu cases
library(tidyterra)
usutu_sp_act<-ggplot2::ggplot() + tidyterra::geom_spatraster(data=r1) +
    ggplot2::geom_point(data=val_agg, ggplot2::aes(x=x, y=y, colour=usutuF)) +
    ggplot2::scale_colour_manual(name="Usutu area", values=val_colours) +
    tidyterra::scale_fill_whitebox_c(name="Density of Usutu cases", palette="arid")+
    ggplot2::theme(aspect.ratio=1,axis.text.x=element_blank(),
                   axis.text.y=element_blank(),axis.ticks=element_blank(),
                   axis.title.x=element_blank(),
                   axis.title.y=element_blank(),)

#the figure for the blackbird part
#
##png("Fig2.png", width=3000, height=3000, pointsize=3,res=300)
library(ggpubr)
ggpubr::ggarrange(ggpubr::ggarrange(usutu_histo,
                                    usutu_sp_act, ncol=2,
                                    labels=c("A", "B"), widths = c(0.6, 1)),
                  p_trends, nrow=2, labels=c("", "C"))

## `stat_bin()` using `bins = 30`.  Pick better value with `binwidth`.

##dev.off()
```

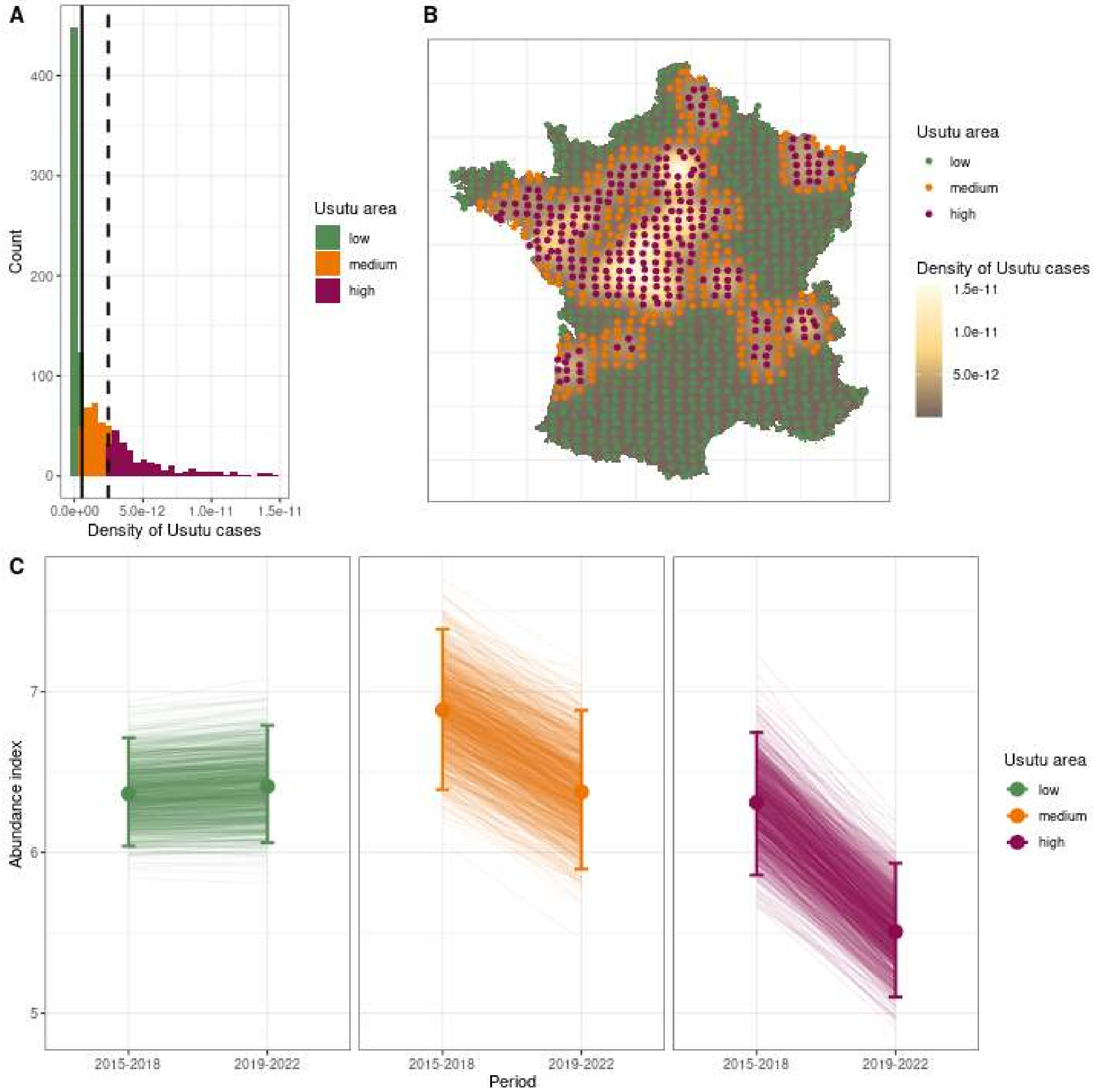
